# An oomycete RXLR effector triggers antagonistic plant hormone crosstalk to suppress host immunity

**DOI:** 10.1101/561605

**Authors:** Ryan Anderson, Devdutta Deb, John Withers, Sheng Yang He, John McDowell

## Abstract

Understanding the mechanisms through which pathogens alter plant cell networks is essential for understanding plant-pathogen interactions and will inform efforts to reduce crop diseases. Oomycetes secrete diverse effector proteins into plant cells. The mechanisms through which these effectors promote virulence are largely unknown. We show that the HaRxL10 effector protein from the *Arabidopsis thaliana* pathogen *Hyaloperonospora arabidopsidis* (*Hpa*) targets a transcriptional repressor (JAZ3) involved in jasmonic acid (JA) signalling. This manipulation activates a regulatory cascade that inhibits salicylic acid (SA) signalling, which normally restricts *Hpa* infection. This virulence mechanism is functionally equivalent to but mechanistically distinct from activation of the antagonistic JA-SA hormone crosstalk by the bacterial JA-mimicking toxin coronatine and by bacterial Type III effectors. These results reveal a key role for JAZ3 in plant immunity and emphasize that JA-SA crosstalk is an Achilles’ heel in the plant immune system, vulnerable to manipulation by diverse microbes.

## INTRODUCTION

Oomycetes are filamentous microbes that resemble fungi but reside within the Chromalveolata (Beakes, Glockling, & Sekimoto, 2012). Many oomycetes cause destructive plant diseases (Kamoun et al., 2015). The majority of plant-pathogenic oomycetes reside within the order *Peronosporales* that includes the genus *Phytophthora* (“plant destroyer”) along with 21 genera of downy mildew pathogens, comprised of over 600 species that are estimated to parasitize at least 15% of flowering plant species (Clark & Spencer-Phillips, 2000; Thines, 2014). The downy mildew pathogen of *Arabidopsis thaliana* (Arabidopsis hereafter) is called *Hyaloperonospora arabidopsidis* (*Hpa* hereafter). This pathogen is frequently found in natural Arabidopsis populations (Holub & Beynon, 1996). Because *Hpa* is one of only two naturally occurring oomycete pathogens of Arabidopsis, it has been adopted as a reference organism for plant-oomycete interactions (McDowell, 2014). *Hpa* is also a reference organism for the obligate biotrophic lifestyle, in which pathogens extract nutrients exclusively from living host cells and cannot survive apart from their hosts (Coates & Beynon, 2010). In keeping with its obligate lifestyle, the *Hpa* genome is configured for stealth and defence suppression in the plant host (Baxter et al., 2010).

A major virulence mechanism, employed by oomycetes and other plant pathogens, is to secrete effector proteins to the exterior (i.e. apoplast) and interior of plant cells (Toruno, Stergiopoulos, & Coaker, 2016). The oomycete secretome can be grouped into several families based on protein sequence motifs (R. H. Jiang & Tyler, 2012). The best-studied family of oomycete effectors is named “RXLR” in reference to a conserved amino acid motif near the N-terminus (Rehmany et al., 2005). The genomes of *Phytophthora* species and downy mildew pathogens contain from approximately 100 to over 1000 genes that are predicted to encode RXLR effector proteins (R. H. Jiang & Tyler, 2012). These proteins are secreted to the interior of plant cells to promote pathogen virulence (Wang et al., 2017). Some RXLR proteins have been shown to interact with plant proteins that regulate the immune signaling network, thereby attenuating the plant’s immune response (Anderson, Deb, Fedkenheuer, & McDowell, 2015). Thus, RXLR proteins are thought to serve as important weapons for *Phytophthora* and downy mildew pathogens. However, much remains to be learned about the mechanism through which RXLR proteins promote virulence.

Like all phytopathogens, oomycetes are highly vulnerable to plant immune responses that are activated when plant surveillance proteins perceive pathogen-derived signals (Jones & Dangl, 2006). For example, plants can recognize a variety of microbe- or pathogen-associated molecular patterns (MAMPs or PAMPS) that are produced as the pathogen multiplies in the apoplast between cells. PAMPs are typically derived from molecules that are broadly conserved between pathogen genera and can be important for pathogen fitness (Boller & Felix, 2009). Often, PAMPs are recognized in the apoplast by transmembrane proteins, called pattern-recognition receptors (PRRs), that engage intracellular signaling cascades to activate PRR-triggered immunity or PTI (Win et al., 2012). Several PAMPs and PRRs have been identified from oomycetes, and oomycete RXLR effectors have been shown to suppress PTI (Raaymakers & Van den Ackerveken, 2016). PTI likely underpins so-called “basal resistance” that limits the growth of virulent oomycetes (Chaparro-Garcia et al., 2011; Ton, Van Pelt, Van Loon, & Pieterse, 2002).

Plants can also recognize pathogen effector proteins that are secreted to the interior of plant cells (Cui, Tsuda, & Parker, 2015). This surveillance is provided by large families of plant NLR proteins that contain a nucleotide binding site and LRRs. NLR proteins often guard plant proteins that are targeted by pathogen effectors to promote virulence and are activated when the guarded protein (or a decoy thereof) is altered by the corresponding effector (Su, Spears, Kim, & Gassmann, 2018). In some instances, NLR proteins can be activated by direct interaction with the corresponding effector (Dangl & McDowell, 2006). Both of these modes of NLR activation can trigger a very fast and potent immune response, termed NLR-triggered immunity or NTI (Win et al., 2012)). For oomycetes, every instance of NTI that has been characterized at the molecular level involves direct or indirect interaction between a plant NLR protein and an oomycete RXLR protein (Anderson et al., 2015). However, oomycetes often co-evolve rapidly to break NTI, and genomic and molecular investigation have revealed that this occurs through a variety of mechanisms (Anderson et al., 2015). Thus, new approaches for developing host resistance would strengthen efforts to mitigate crop damage caused by oomycetes.

PTI and NTI are regulated through a complex, highly interconnected network of signaling sectors that provide strength and robustness (Katagiri & Tsuda, 2010; Nobori, Mine, & Tsuda, 2018). The two best-studied immune signaling sectors are mediated by the oxylipin phytohormone jasmonic acid (JA) and the phenolic phytohormone salicylic acid (SA) (Klessig, Choi, & Dempsey, 2018; Pieterse, Leon-Reyes, Van der Ent, & Van Wees, 2009). JA responses are induced by chewing insects and by necrotrophic pathogens that kill plant cells (Figure 1-figure supplement 1A) (Glazebrook, 2005; Tesfaye Mengiste, 2011). SA signalling is typically activated by biotrophic pathogens that extract nutrients from living plant cells (Grant & Jones, 2009; Katagiri & Tsuda, 2010). Importantly, the JA and SA signalling sectors can be mutually antagonistic, such that activation of one sector can inhibit the other (Figure 1-figure supplement 1A) (Kemal Kazan & Lyons, 2014). Although exceptions to the above generalities have been documented, JA-SA cross-talk is widespread in plants and likely evolved before the angiosperm-gymnosperm divergence (Han, 2017; Thaler, Humphrey, & Whiteman, 2012). JA-SA antagonism is thought to provide optimal defence, through which immune responses can be tailored to specific types of pathogens and thereby reduce costs of resistance (Spoel, Johnson, & Dong, 2007). However, JA-SA antagonism also provides a mechanism for pathogens to suppress one sector by inducing the other, thereby activating ineffective responses while suppressing those that would otherwise limit growth of the invader (Grant & Jones, 2009).

**Figure 1.**
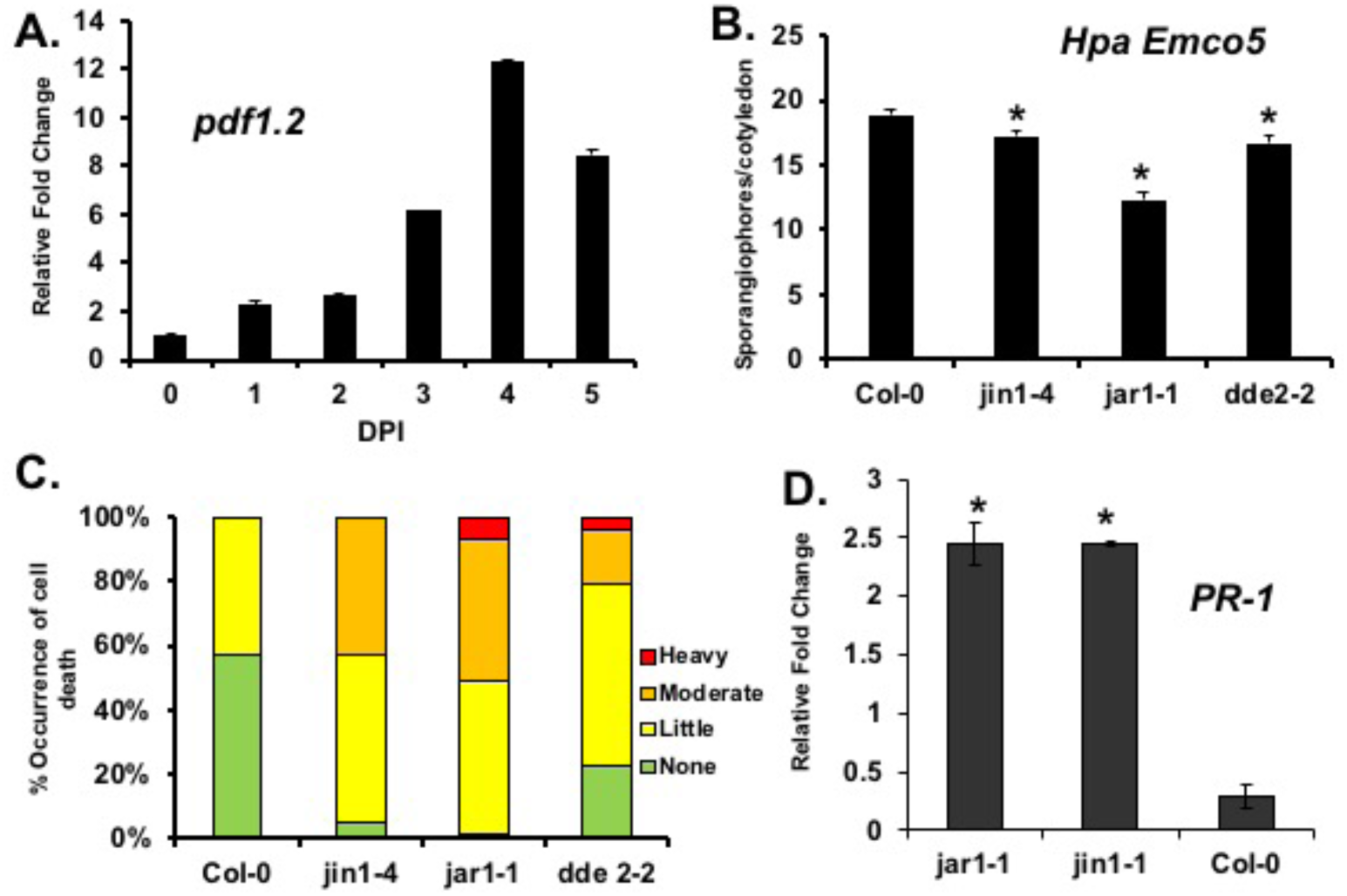
Evidence that *Hpa* engages the *Arabidopsis* jasmonic acid signalling sector to suppress SA-mediated immunity. (**A**) Elevated transcription of JA marker gene *pdf1.2* during infection of Arabidopsis Col-0 by virulent *Hpa* Emco5. Quantitative, reverse-transcriptase PCR (qRT-PCR) was used to measure transcript abundance of *pdf1.2* at the indicated days post infection (DPI) Transcripts were normalized to *AtACTIN2*, and fold change was calculated relative to 0 DPI. Error bars represent standard deviation among technical replicates. (**B**) Reproduction of virulent *Hpa* Emco5 in JA biosynthesis (*dde2* and *jar1*) and signaling (*jin1*) mutants. Col-0 and JA mutant plants were challenged with virulent *Hpa* Emco5. Disease progression was quantified 7 DPI by visual sporangiophore counts. Error bars depict standard error of the mean (SEM). (**C**) Host cell death in response to growth of virulent *Hpa* Emco5 in JA mutants, based on visual assessment of trypan blue-stained samples from the indicated *Hpa* Emco5-infected plants. (**D**) Elevated transcript abundance of the SA marker gene *PR-1* in *jar1* and *jin1* mutants (note that *dde2* was not tested). mRNA was extracted from uninfected plants, and transcript abundance was measured using qRT-PCR. *PR-1* transcript abundance was normalized to *AtACTIN2*. Asterisks designate statistically significant (p < 0.05) differences between mutants and Col-0 plants. All experiments were repeated at least three times with similar results.

Jasmonic acid signalling impacts a variety of plant physiological responses in addition to immune responses. These include plant growth, development and responses abiotic stresses (Howe, Chung, Niu, & Browse, 2009). JAZ proteins (Jasmonate, Zim Domain) are co-receptors for JA and key regulators of JA-dependent responses (Pauwels & Goossens, 2011). JAZ proteins repress JA-responsive genes under conditions in which JA responses are not induced (Browse, 2009). This repression involves binding and inhibition of transcription factors (e.g., MYC2, MYC3, MYC4) that act as positive regulators of JA-responsive genes (Figure 1-figure supplement 1B), (K. Kazan & Manners, 2013). This repression is relieved when pathogens, insects, or developmental signals induce biosynthesis of JA and its bioactive form, JA-isoleucine (Browse, 2009). JA-Ile binds to and activates a Skp/Cullin/F box-type ubiquitin ligase complex, SCF^COI1^, in which Coronatine-insensitive 1 (COI1) is the F-box protein (Fonseca, Chico, & Solano, 2009). JAZ proteins are recruited to SCF^COI1^ via JA-Ile binding, and are subsequently ubiquitinated for destruction in the 26S proteasome, thereby releasing JA-responsive transcriptional activators from repression (Browse, 2009). This mechanism is evolutionarily conserved in land plants (Han, 2017). *Arabidopsis* encodes a family of 12 JAZ proteins. The majority of *At*JAZ proteins appear to be regulated by SCF^COI1^ (Browse, 2009). Some of the *At*JAZ proteins interact with each other, and there is redundancy and specificity in the proteins/genes that are regulated by individual JAZ family members (Chini, Gimenez-Ibanez, Goossens, & Solano, 2016). As such, SCF^COI1^, along with the JAZ proteins and their collective common and unique targets, represent a highly connected regulatory hub that influences many important plant processes. However, the mechanisms through which the SCF^COI1^-JAZ hub regulates diverse downstream responses are incompletely understood (Chini et al., 2016).

Evidence for manipulation of JA-SA antagonism has accumulated for diverse pathogens and pests, as well as mutualists, in aerial organs and in roots (Gutjahr & Paszkowski, 2009; Zhang, Zhang, Melotto, Yao, & He, 2017). The mechanisms underlying this manipulation are now beginning to emerge, led by research on the bacterial pathogen *Pseudomonas syringae*: Some *P. syringae* strains secrete the small molecule coronatine (COR) that acts as a molecular mimic of JA and promotes virulence in part by triggering degradation of JAZ proteins, thereby activating a molecular cascade that suppresses SA signalling (Brooks, Bender, & Kunkel, 2005; Du et al., 2014; Gimenez-Ibanez et al., 2017; Katsir, Schilmiller, Staswick, He, & Howe, 2008; Melotto et al., 2008; Zheng et al., 2012). Moreover, *P. syringae* can deploy Type III effector proteins that degrade most or all of the JAZ proteins in Arabidopsis (Gimenez-Ibanez et al., 2014; S. Jiang et al., 2013). For fungi, an effector from the mutualist *Laccaria bicolor* targets a poplar JAZ protein to promote root colonization, but this virulence mechanism does not appear to involve SA-JA cross talk (Plett et al., 2014). For oomycetes, an *Hpa* effector targets the mediator transcriptional regulatory complex, downstream of the SCF^COI1^-JAZ module, to impinge on JA-SA crosstalk (Caillaud et al., 2013). In this manuscript, we describe evidence that an *Hpa* RLXR protein manipulates JA-SA antagonism to suppress SA-mediated immunity. This work reveals a specific role for the JAZ3 protein as a regulator of immunity and extends recent observations that the JA receptor complex is exploited as an Achilles Heel by diverse microbes.

## Results

*Genetic Evidence that JA signalling is relevant to Hpa virulence:* The research in this report followed from our unexpected observation that the JA marker gene *pdf1.2* is induced during infection by a virulent isolate of *Hpa* (Fig. 1A). This was surprising because previously published experiments have demonstrated that SA-mediated responses are important in *Arabidopsis* for immunity against *Hpa*, while JA-mediated responses do not contribute to resistance against *Hpa* (Glazebrook, 2005; McDowell et al., 2000; Thomma et al., 1998). Thus, it seemed unlikely that the plant host was activating JA signalling as an immune response against *Hpa*. Based on precedent provided by studies of coronatine, we hypothesized that *Hpa* could activate JA signalling to suppress SA-mediated responses. As an initial challenge of this hypothesis, we tested the *jar1*, *jin1*, and *dde2* mutants that compromise the function of the JA signalling sector (Figure 1-figure supplement 1A)(Browse, 2009). We reasoned that if the JA signalling sector is co-opted by *Hpa* to suppress SA signalling, then Arabidopsis mutants disabling the JA sector should display reduced susceptibility to *Hpa*, accompanied by elevated SA signalling. Accordingly, we observed that the *jar1* mutant displayed reduced sporulation by a virulent *Hpa* isolate (Fig. 1B). JAR1 encodes an enzyme that conjugates JA to Ile (Staswick & Tiryaki, 2004); thus, biosynthesis of JA-Ile is genetically essential for full *Hpa* virulence, similar to bacteria (Laurie-Berry, Joardar, Street, & Kunkel, 2006). A similar but weaker phenotype was exhibited by the *jin1* mutant that affects the JA-responsive transcription factor MYC2 (K. Kazan & Manners, 2013) and the JA biosynthesis mutant *dde2* (von Malek, van der Graaff, Schneitz, & Keller, 2002). All three mutants displayed enhanced plant cell death around *Hpa* infection structures, compared to the wild type Col-0 (Fig. 1C). *jar1* and *jin1* exhibited elevated expression of the SA marker gene *PR-1* (Fig. 1D), suggesting that removal of JA signalling in these mutants leads to de-repression of SA-mediated immunity. These results establish that, by genetic criteria, the JA signaling sector is necessary for full *Hpa* virulence and negatively regulates host immune responses, consistent with the hypothesis that *Hpa* manipulates JA signaling to promote virulence.

*The AtPPIN1 database identifies a link between an Hpa RXLR protein and a negative regulator of JA-responsive genes:* To identify potential mechanisms through which *Hpa* could manipulate JA signalling, we examined the *Arabidopsis* Plant-Pathogen Interactome database (*At*PPIN1) that documents putative targets of *Hpa* RXLR effectors identified by yeast two-hybrid screens (Mukhtar et al., 2011). We examined the *At*PPIN1 dataset specifically for interactions between *Hpa* RXLR proteins and plant proteins that are linked to the JA signalling sector. We found only one direct connection, involving the *Hpa* effector HaRxL10 and the JA transcriptional repressor JAZ3 (Figure 2-figure supplement 1A). The *At*PPIN1 data predict that HaRxL10 interacts with JAZ3, along with four other transcription factors, a subunit of the COP9 complex, CSN5A, and an E3 ligase protein, BOI1 (Figure 2-figure supplement 1B). A previous publication reported that a *jaz3* knockout mutant displays enhanced susceptibility to virulent *Hpa* (Mukhtar et al., 2011). We reproduced this result using assays for pathogen fitness (sporulation, Fig. 2A) and growth *in planta* (measured by qPCR, Fig. 2B). The enhanced growth and fitness of *Hpa* in a *jaz3* null mutant indicates that the HaRxL10-JAZ3 interaction is biologically relevant for *Hpa* virulence.

**Figure 2.**
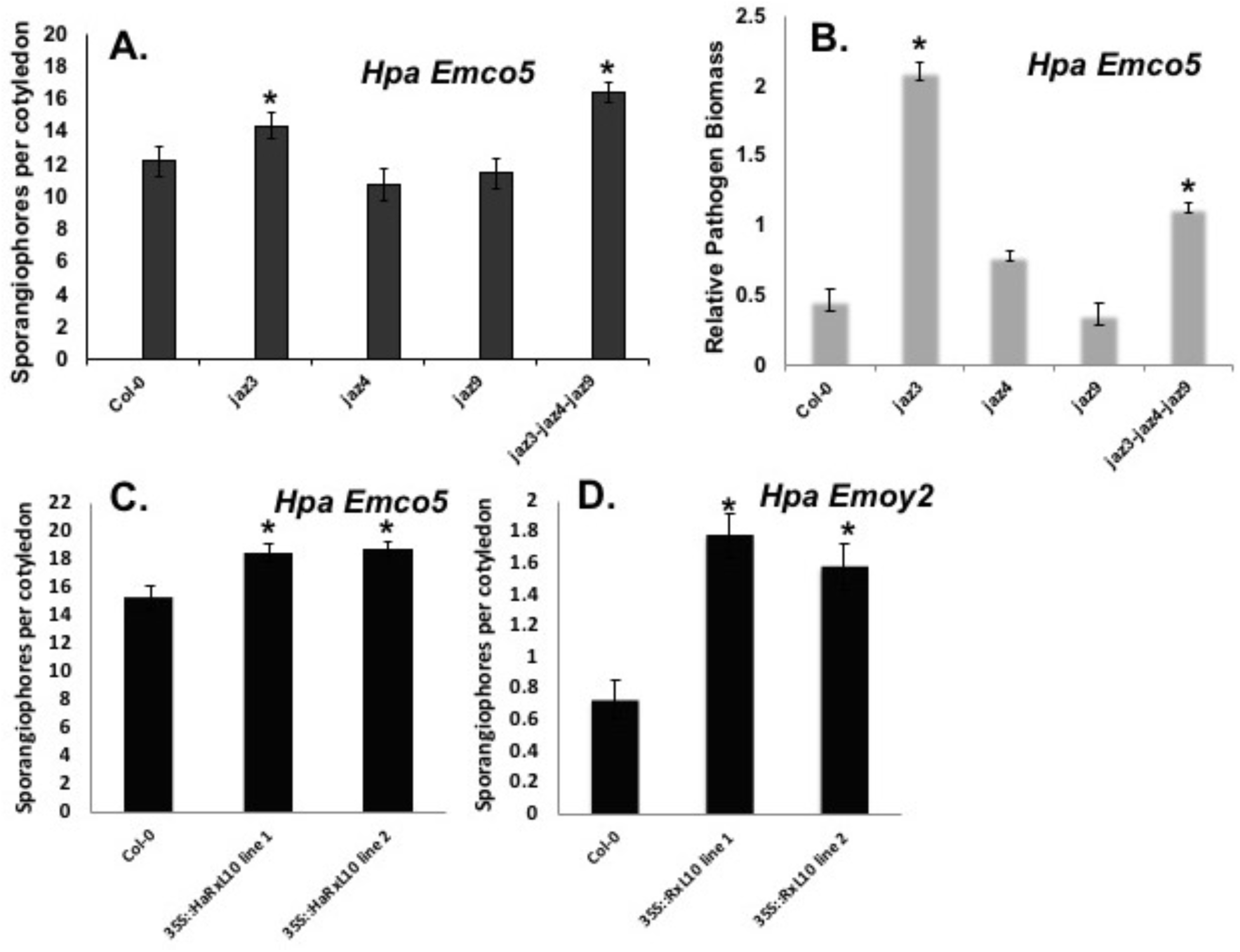
Susceptibility to *Hpa* is enhanced in a *jaz3* loss-of-function mutant and in transgenic plants that overexpress HaRxL10. (**A**) Reproduction of virulent *Hpa* Emco5 in *jaz3, jaz4*, *jaz9* single mutants and a j*az3-jaz4-jaz9* triple mutant, based on visual sporangiophore counts. (**B**) Hyphal growth of virulent *Hp*a Emco5 quantified via PCR in the *jaz* mutants and control Col-0 plants. Genomic DNA was extracted from seedlings collected at six days post inoculation. qPCR was used to measure the relative abundance of *HpaACTIN* relative to *AtACTIN2*, as a proxy for pathogen biomass. Error bars represent SE of technical replicates. (**C, D**) Reproduction of virulent *Hpa* Emco5 (**C**) or avirulent *Hpa* Emoy2 (**D**) on *Arabidopsis* Col-0 lines stably transformed with a CaMV35S-RxL10 transgene. Line 1 and Line 2 are independent transgenic lines. * P < 0.05; t-test comparisons with Col-0. All experiments were repeated at least three times with similar results.

*HaRxL10 is transcribed during infection and promotes Hpa virulence:* The predicted HaRxL10 protein contains a signal peptide followed by RXLR and EER motifs and a C-terminal domain with no recognizable motifs (Figure 2-figure supplement 1C). This effector protein has not been functionally characterized, to our knowledge. We determined that the *HaRxL10* gene is induced *in planta* during the early stages of *Hpa* infection, consistent with a virulence function (Figure 2-figure supplement 1D). We constructed transgenic Arabidopsis Col-0 containing a transgene expressing the predicted processed form of HaRxL10, driven by the CaMV35S promoter. Reproduction of two *Hpa* isolates was enhanced in these lines (Fig. 2C, D), indicating that HaRxL10 can function as a virulence-promoting protein.

*JAZ3 interacts with HaRxL10 in yeast and is genetically essential for basal resistance to Hpa:* In directed yeast two-hybrid assays we confirmed that HaRxL10 interacts with JAZ3 in yeast (Fig. 3A). HaRxL10 also interacts in yeast with JAZ4 and JAZ9 but not JAZ1, 2, 6, 10, or 12 (Fig. 3A). RxL10 and COI1 do not interact under these conditions (Fig. 3A). Yeast two-hybrid experiments with deletion mutants of JAZ9 indicate that the N-terminal domain is important for the interaction with HaRxL10, while the ZIM and JAZ domains are dispensable (Fig. 3B).

**Figure 3.**
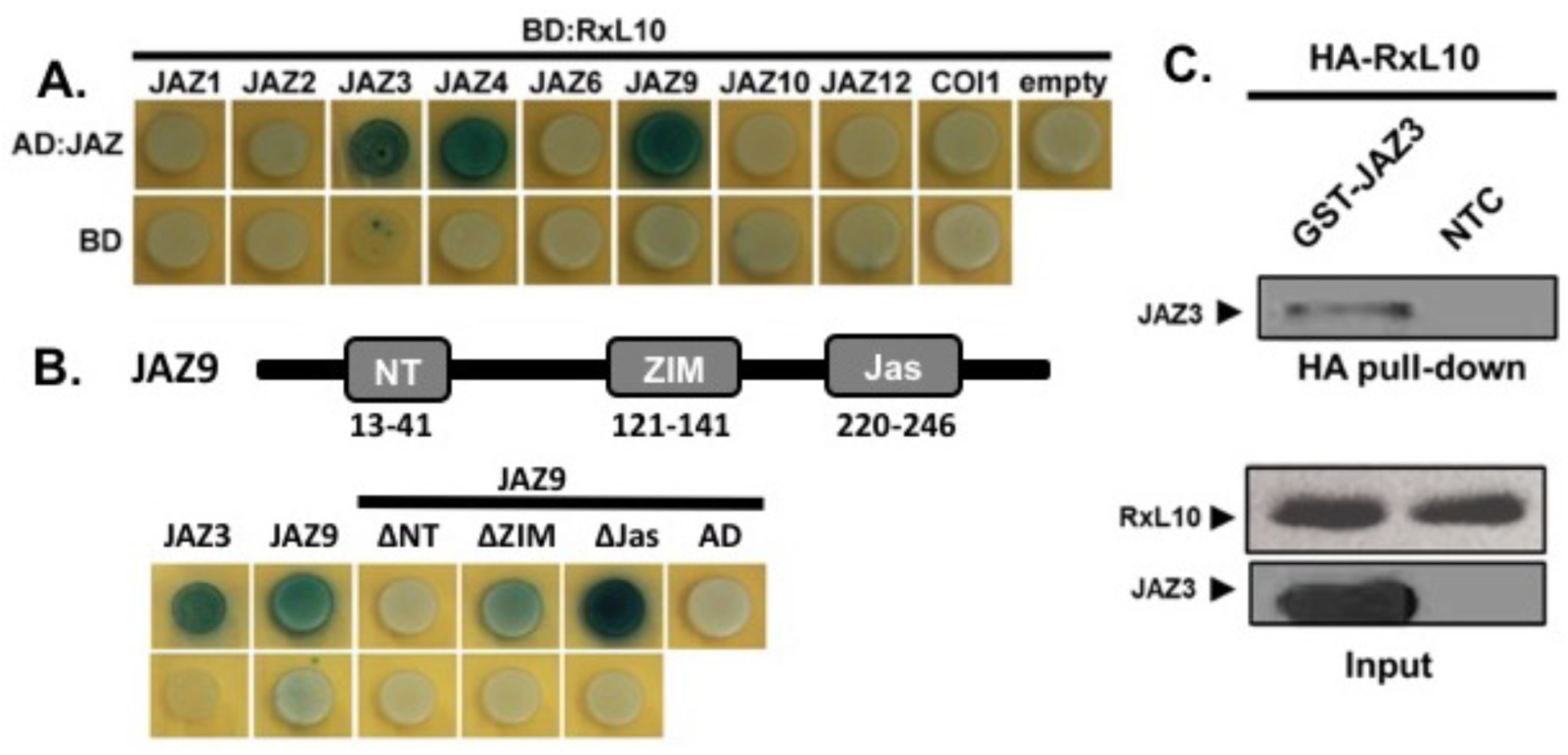
Interaction between HaRxL10 and Arabidopsis JAZ proteins **(A)** Yeast two-hybrid (Y2H) assays for interaction between HaRxL10 and *Arabidopsis* JAZ proteins. AD and BD refers to control constructs containing activation and DNA binding domains of the LexA-based Y2H system, respectively. **(B)** Y2H assays for interaction between HaRxL10 and deletion derivatives of the *Arabidopsis* JAZ9 protein. The schematic of the JAZ9 protein depicts functional domains that were deleted**. (C)** I*n-vitro* co-immunoprecipitation assay (co-IP) for interaction between HaRxL10 and *Arabidopsis* JAZ3. HA-HaRxL10, GST-JAZ3 or NTC (no template control) were synthesized *in vitro* using the TNT^®^ Coupled Wheat Germ Extract Systems (Promega). Cell lysates were then immunoprecipitated using anti-HA antibody. The immunoprecipitates were examined by Western blotting using anti-GST antibody. Input represented 10% of wheat germ lysates used in the co-IP experiment. Co-IP experiments using GST pull-downs also demonstrated interaction (not shown). All experiments were repeated at least three times with similar results.

Because HaRxL10 interacts with JAZ4 and JAZ9, we tested the effect of previously validated loss-of-function mutants of these genes to determine their role during infection. We challenged *jaz4* and *jaz9* mutants, along with a *jaz3*/*jaz4*/*jaz9* triple mutant, with virulent *Hpa*. We observed no enhanced virulence of *Hpa* in the *jaz4* and *jaz9* single mutants (Fig. 2A, B). The triple mutant supported enhanced *Hpa* sporulation relative to wild-type Col-0, but the amount of sporulation was not significantly enhanced compared to the *jaz3* single mutant (Fig. 2A). Vegetative growth in the triple mutant varied but was generally equivalent or less than the amount of growth supported in the *jaz3* single mutant. These results indicate little or no effect from the *jaz4* and *jaz9* mutants (Fig. 2B). Thus, by genetic criteria, *JAZ3* plays a major role in basal resistance to *Hpa*, and we focused our subsequent experiments on *JAZ3*.

*JAZ3 interacts with RXL10* in vitro *and in planta*: We confirmed that HaRxL10 interacts with JAZ3 in an *in vitro* co-immunoprecipitation assay, suggesting that the proteins can bind directly to each other in the absence of other plant or oomycete proteins (Fig. 3C). Bimolecular fluorescence complementation experiments in *Nicotiana benthamiana* indicate that HaRxL10 and JAZ3 interact in the nucleus and the cytoplasm (Fig. 4A). Fluorescently tagged HaRxL10 and JAZ3 exhibit a nucleocytoplasmic distribution when expressed individually (Fig. 4B), including a strong signal from subnuclear structures of unknown function (Fig. 4C) that resemble structures previously described for localization of JAZ9 and JAZ3 (Withers et al., 2012; Yang et al., 2017).

**Figure 4.**
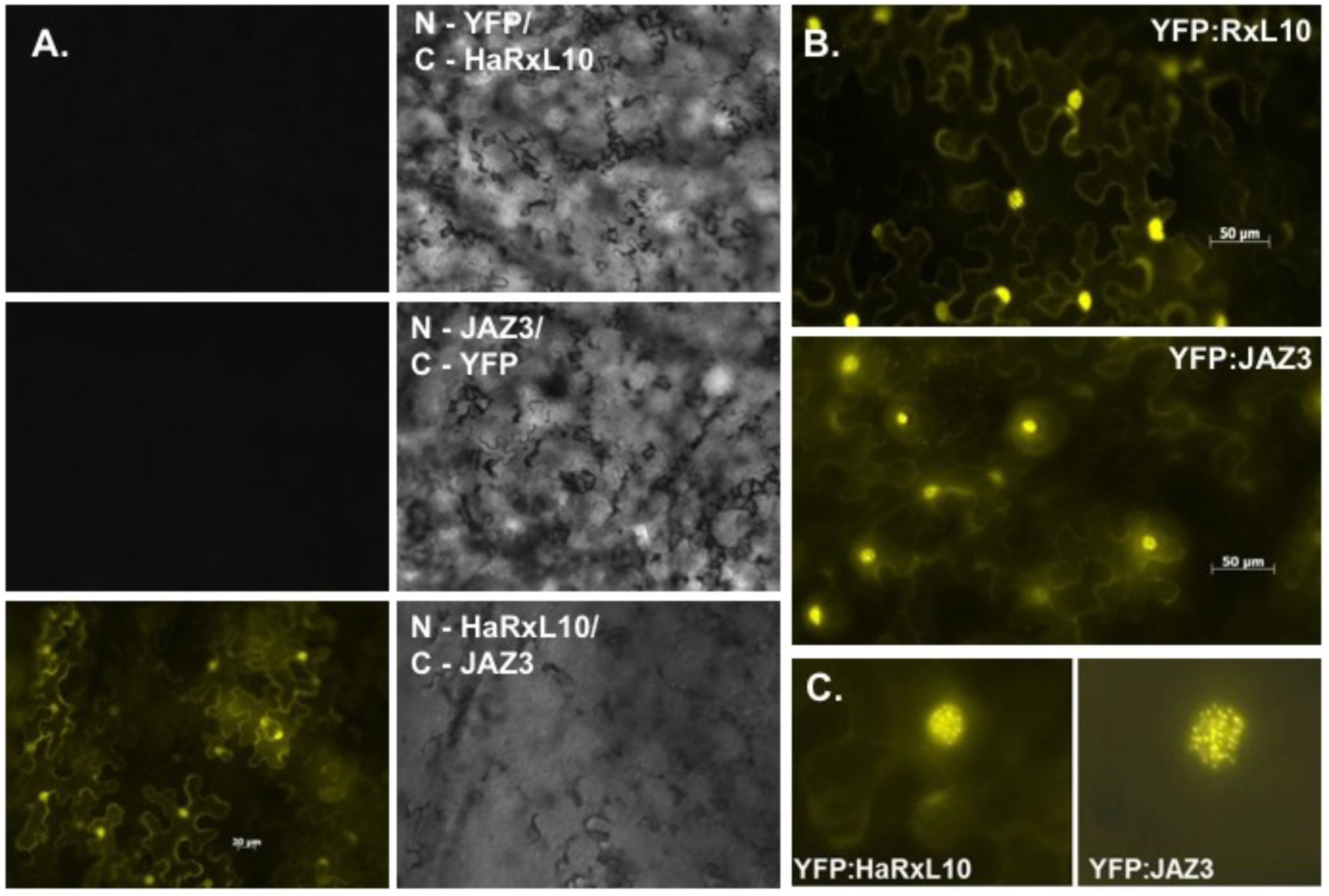
Interaction and subcellular localization of RxL10 and JAZ3 *in planta*. **(A)** BiFC-based protein interaction assay in *Nicotiana benthamiana* of split YFP constructs of JAZ3-YC with an empty N-YFP vector, YN-RxL10 with an empty C-YFP vector and JAZ3-YC with YN-HaRxL10. (**B, C**) Confocal microscopic images of transiently expressed, YFP-tagged HaRxL10 and JAZ3 in *N. benthamiana* epidermal cells. All experiments were repeated at least three times with similar results.

*Genetic evidence that the HaRxL10 effector activates JA signalling:* As described above, bacterial coronatine (COR) is a molecular mimic of JA that triggers degradation of JAZ proteins and activates a signalling cascade that suppresses SA-dependent immune responses (Melotto, Underwood, Koczan, Nomura, & He, 2006; Zheng et al., 2012). Importantly, COR antagonizes stomatal defence, in which the plant closes stomata to impede the bacteria’s ability to travel from the leaf surface to the interior (Melotto, Zhang, Oblessuc, & He, 2017). To test whether HaRxL10 affects stomatal defence in a manner similar to COR, we transformed a *P. syringae* COR biosynthetic mutant (*Pst* DC3118, cor-(Brooks et al., 2004)) with a plasmid configured to express HaRxL10 fused with the N-terminal leader of the bacterial AvrRps4 protein that can be translocated into plant cells through the Type III secretion system (Fig. 5A, (Sohn, Lei, Nemri, & Jones, 2007)). This approach was previously used to test for functional equivalence between COR and *P. syringae* Type III effectors (Gimenez-Ibanez et al., 2014; S. Jiang et al., 2013). We inoculated Arabidopsis plants by spraying the leaf surface with bacteria, and then assessed plant disease symptoms and measured bacterial growth in the interior of the leaf at three days post-infection. In this assay, translocated HaRxL10 partially rescued the virulence defects (*in planta* growth and disease symptoms) of the cor^−^ mutant when bacteria were sprayed onto the leaf surface (Fig. 5B-C). The invasion/growth defect of the cor^−^ strain was also partially rescued when HaRxL10 was expressed from a plant transgene in stably transformed Arabidopsis Columbia (Col-0) plants (Fig. 5D). Similar to COR, HaRxL10 did not enhance virulence when bacteria were inoculated directly into the apoplast by syringe infiltration, indicating that the virulence-promoting effect in the spray assay was due to interference with stomatal defence. In comparison, HaRxL10 did not enhance the virulence of wild-type *Pst* DC3000 and only slightly increased the virulence of the *Pst* DC3000 Δ*CEL* mutant (Fig. 5E), which contains a genomic deletion that removes three Type III effectors in the conserved effector locus (CEL) that promote virulence through mechanisms that differ from that of COR (DebRoy, Thilmony, Kwack, Nomura, & He, 2004).

**Figure 5.**
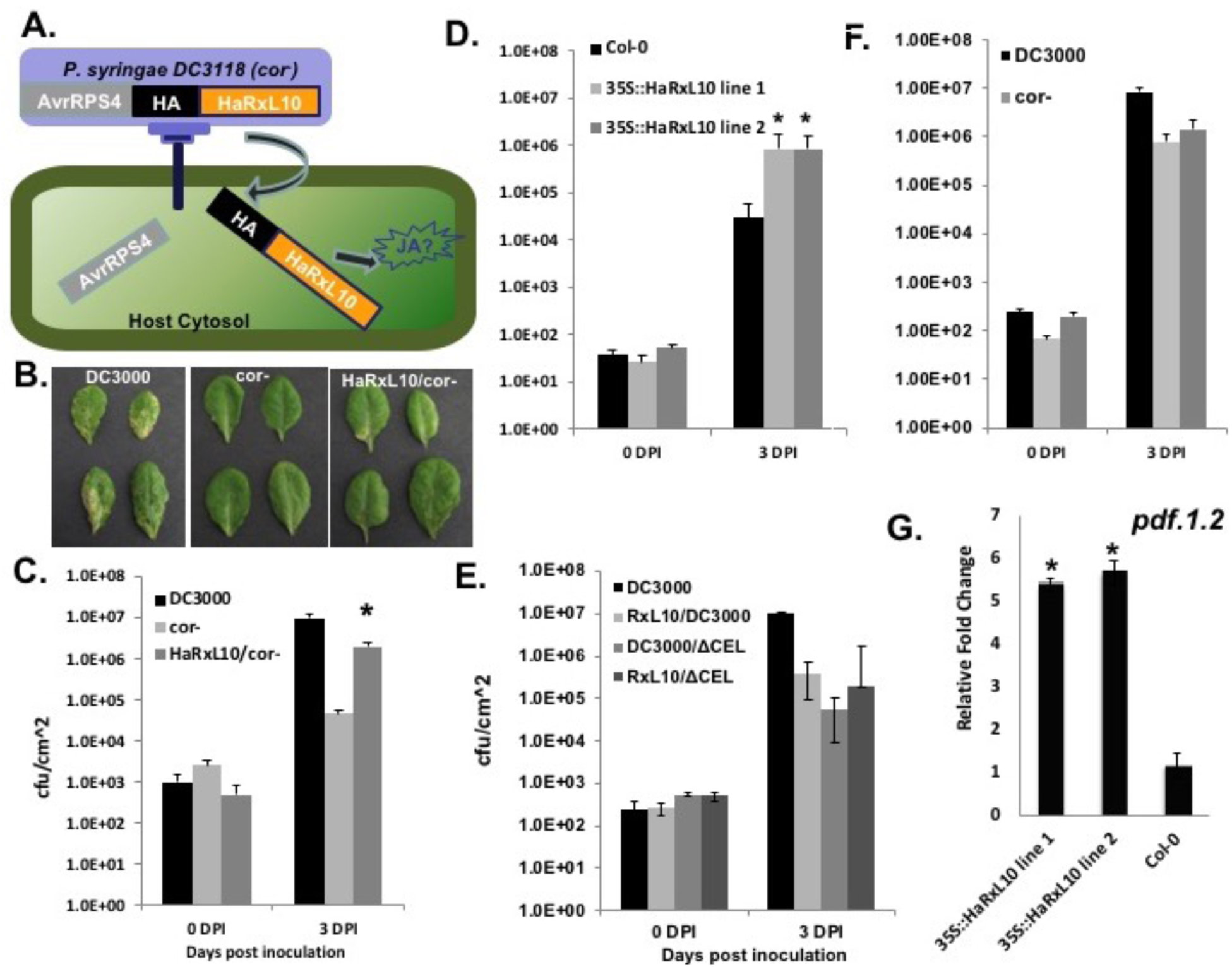
Genetic evidence that HaRxL10 targets the JA pathway. (**A**) Schematic of the effector detector system through which a fusion of HaRxL10 and the bacterial AvrRps4 leader is delivered from *P. syringae* via Type III secretion. (**B**) Plant disease symptoms triggered by *Pst* DC3000 and the cor^−^ mutant Pst DC3118, with or without pEDV-HaRxL10. (**C**) *In planta* growth, following spray inoculation of Col-0, of *Pst* DC3118 with or without HaRxL10. (**D**) *In planta* growth of *Pst* DC3118, without HaRxL10, on *Arabidopsis* Col:35S-RXL10 transgenic lines. (**E**) *In planta* growth of *Pst* DC3000 and *Pst* DC3000/ΔCEL with or without HaRxL10. (**F**) *In planta* growth, following spray inoculation of Col-0, of *Pst* DC3118 with or without HaRxL10. (**G**) Transcript abundance of the JA marker gene *pdf1.2* in uninfected Col:35S-RxL10 plants, assayed by qRT-PCR. All experiments were repeated at least three times with similar results.

Moreover, HaRxL10 did not promote virulence of the cor^−^ mutant when bacteria were injected into the leaf apoplast, as was previously demonstrated for COR (Fig. 5F). This specificity suggests that the mechanism by which HaRXL10 promotes virulence is more similar to that of COR than to the Type III effectors encoded in the CEL. Additionally, the JA marker gene *pdf1.2* is constitutively induced in uninfected Col:*35S-HaRxL10* (Fig. 5G), demonstrating that transgenic expression of HaRxL10 is sufficient to activate JA responses, even in the absence of pathogen infection. Together, these experiments indicate that the mode of HaRxL10-dependent virulence is similar to that of coronatine.

*RxL10 targets JAZ3 for proteosomal degradation:* Because loss of *JAZ3* is sufficient to enhance susceptibility to virulent *Hpa* (Fig. 2A, B)(Mukhtar et al., 2011), we hypothesized that HaRxL10 degrades or otherwise nullifies JAZ3 to promote virulence. Accordingly, JAZ3-YFP abundance was substantially reduced by co-expression with HaRxL10 in *N. benthamiana*, albeit to a lesser extent than when leaves expressing JAZ3-YFP were treated with 10mM methyl jasmonate (Fig. 6A, B). In Arabidopsis, abundance of transgenically expressed YFP-JAZ3 was reduced during colonization by virulent *Hpa* (Fig. 6C). JAZ3-The HaRxL10-dependent destabilization of JAZ3 in *N. benthamiana* is reversed by the addition of proteasome inhibitor MG132 *in vivo* (Fig. 6D). These data indicate that HaRxL10 targets JAZ3 for degradation by the 26S proteasome.

**Figure 6.**
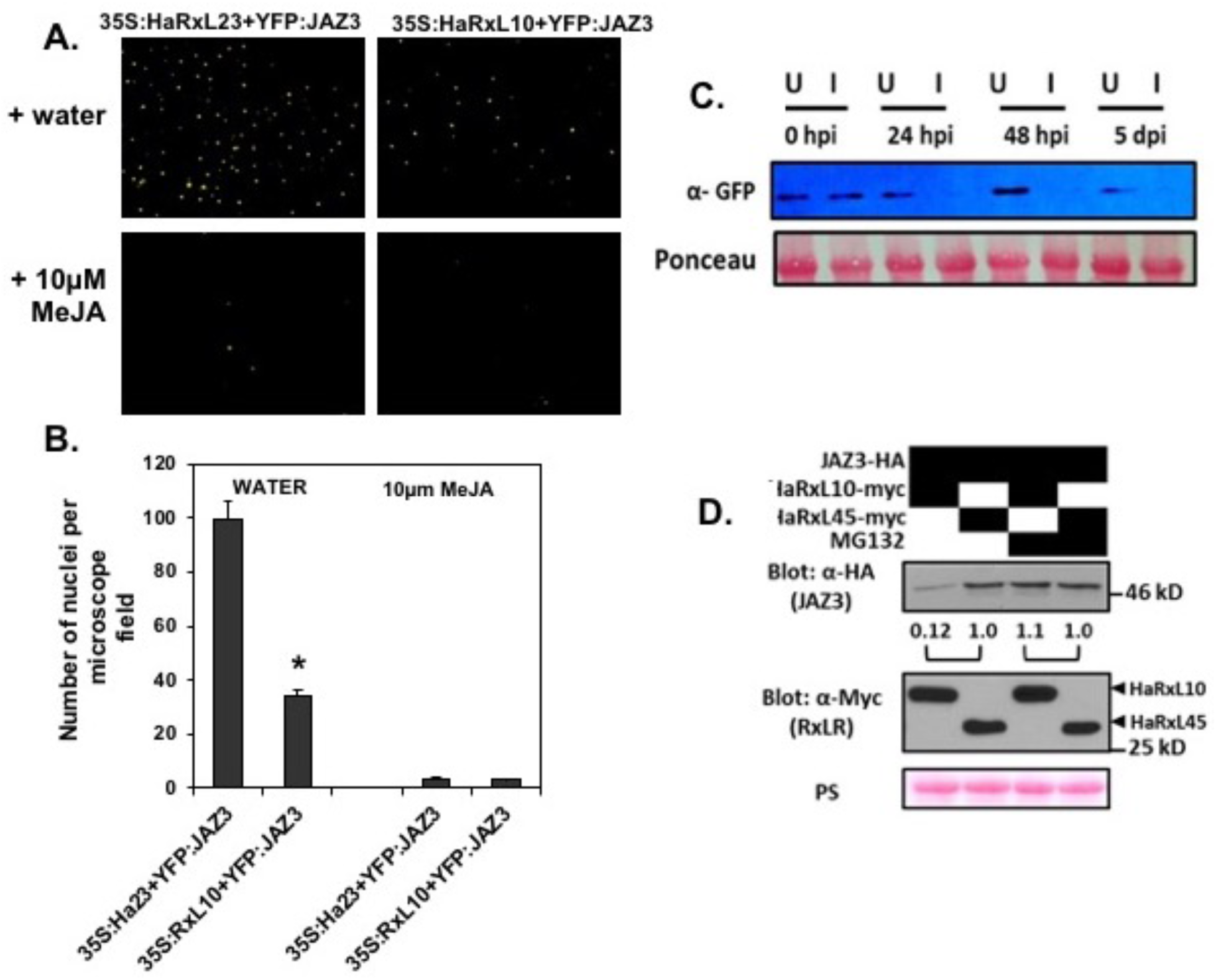
HaRxL10 reduces JAZ3 protein abundance. (**A**) Confocal micrographs depicting reduced signal from YFP-JAZ3 when co-expressed with HaRxL10 in *N. benthamiana* with water or methyl jasmonate (MeJA) treatment. HaRxL23 is a control RXLR effector from *Hpa* that does not impact the JA-SA crosstalk. (**B**) Quantification of YFP-JAZ3 signal from nuclei of *N. benthamiana* epidermal cells. (**C**) Western blot showing reduced abundance of YFP-JAZ3 in transgenic *Arabidopsis* seedlings colonized by virulent *Hpa* Emco5. “U” refers to uninfected and “I” refers to infected seedlings. (**D**) Western blot showing reduced abundance of JAZ3 when co-expressed with HaRxL10 in *N. benthamiana*. HaRxL45 is a control effector that does not interact with JAZ3 or affect SA-JA crosstalk but is localized to subnuclear foci (Yang et al., 2017). The 26S proteasome inhibitor MG132 suppresses HaRxL10-dependent degradation of JAZ3. Numbers depict relative band intensities. All experiments were repeated at least three times with similar results.

*HaRxL10 activates the same downstream signalling cascade as coronatine:* A molecular cascade through which coronatine suppresses *Arabidopsis* SA-mediated defences has been identified (Zheng et al., 2012) (Figure 1-figure supplement 1C): COR mimics JA-Ile to induce SCF^COI1^-dependent degradation of JAZ proteins, thereby derepressing the MYC2/3/4 transcription factors and activating three MYC2-regulated *NAC* (petunia NAM and *Arabidopsis* ATAF1, ATAF2, and CUC2) transcription factor genes: *ANAC019*, *ANAC055*, and *ANAC072*. In turn, the NAC proteins directly repress expression of a key SA biosynthetic gene (isochorismate synthase 1, *ICS1*) and activate genes encoding SA glucosyl transferase gene 1 (*SAGT1*) and SA methyltransferse (*BSMT1*), which metabolize SA and render it biologically inactive. Thus, this genetic reprogramming reduces the pool of bioactive SA and thereby compromises the SA-regulated immune signalling sector. Because HaRxL10 partially compensates for COR deficiency in *Pst* DC3118, we hypothesized that the HaRxL10-JAZ3 interaction activates the same cascade. Accordingly, transcript abundance of *ANAC019*, *ANAC055*, *ANAC072* was elevated in uninfected Col:*35S-HaRxL10* compared to control plants. *SAGT1* transcript abundance was also elevated in uninfected Col:*35S-HaRxL10*, while *ICS1* transcript abundance is reduced (Fig. 7). Finally, basal transcript abundance of PR-1 is reduced in uninfected Col:*35S-HaRxL10*. These data indicate that expression of HaRxL10 is sufficient to activate the same downstream cascade that mediates JA-SA antagonism triggered by COR.

**Figure 7.**
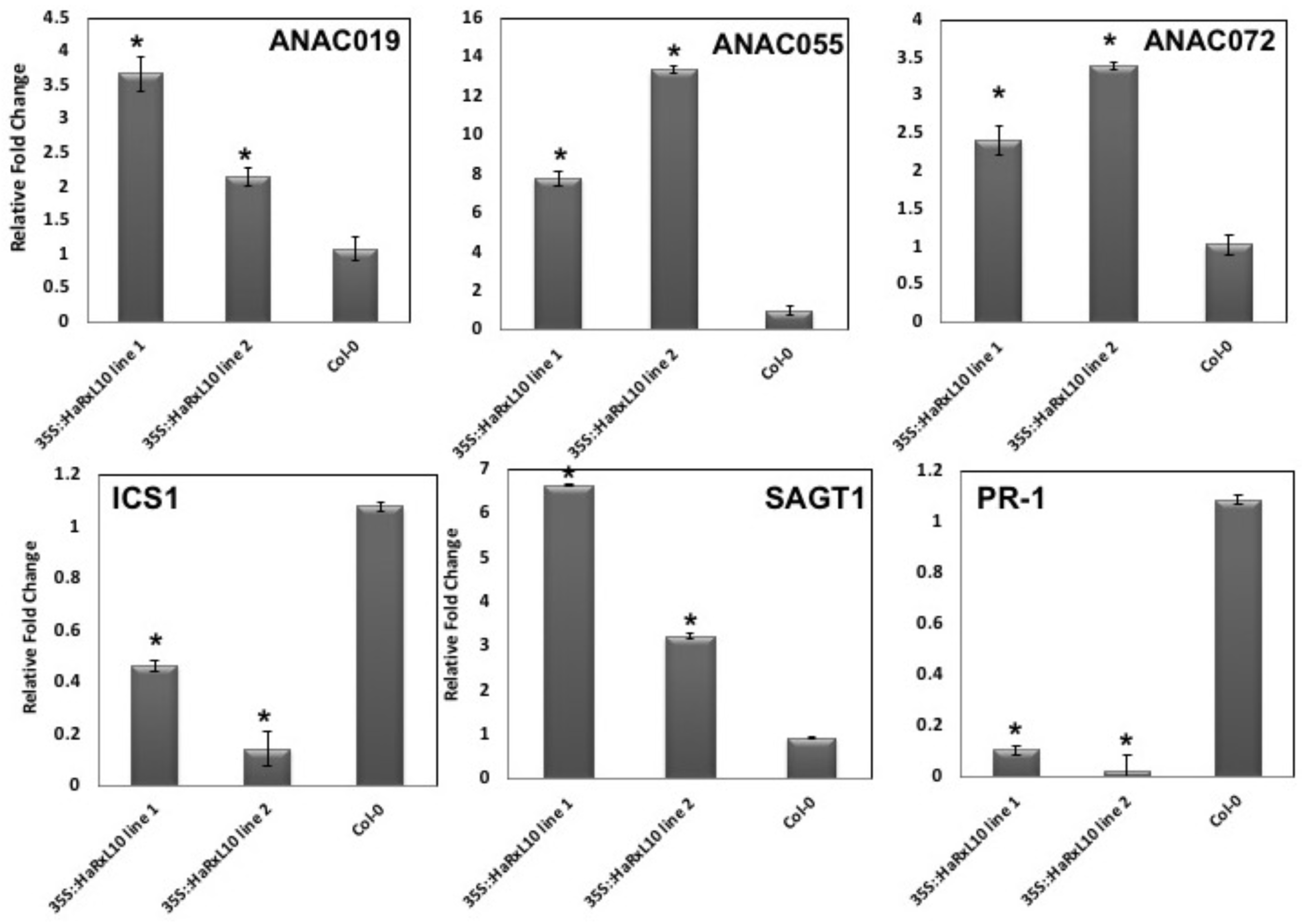
Transgenic expression of HaRxL10 is sufficient to activate a regulatory cascade that quells SA signalling. Transcript abundance was measured using qRT-PCR using cDNA from uninfected Col:*35S-RxL10* OX line 1 and line 2 with primers specific for the indicated genes. Transcript abundance was normalized to *AtACTIN2*. Error bars depict variance among technical replicates. * ddCt values representing statistically significant (\**P* < 0.05) differences between transgenic lines and Col-0. This experiment was repeated three times with similar results.

Similar effects of HaRxL10 overexpression on each of the six genes were seen in infected plants: transcript abundance of *ANAC019*, *ANAC055*, *ANAC072* and *SAGT1* was elevated in Col:*35S-HaRxL10* compared to control plants (Fig. 8), while the abundance of *ICS1* and *PR-1* transcripts were reduced.

**Figure 8.**
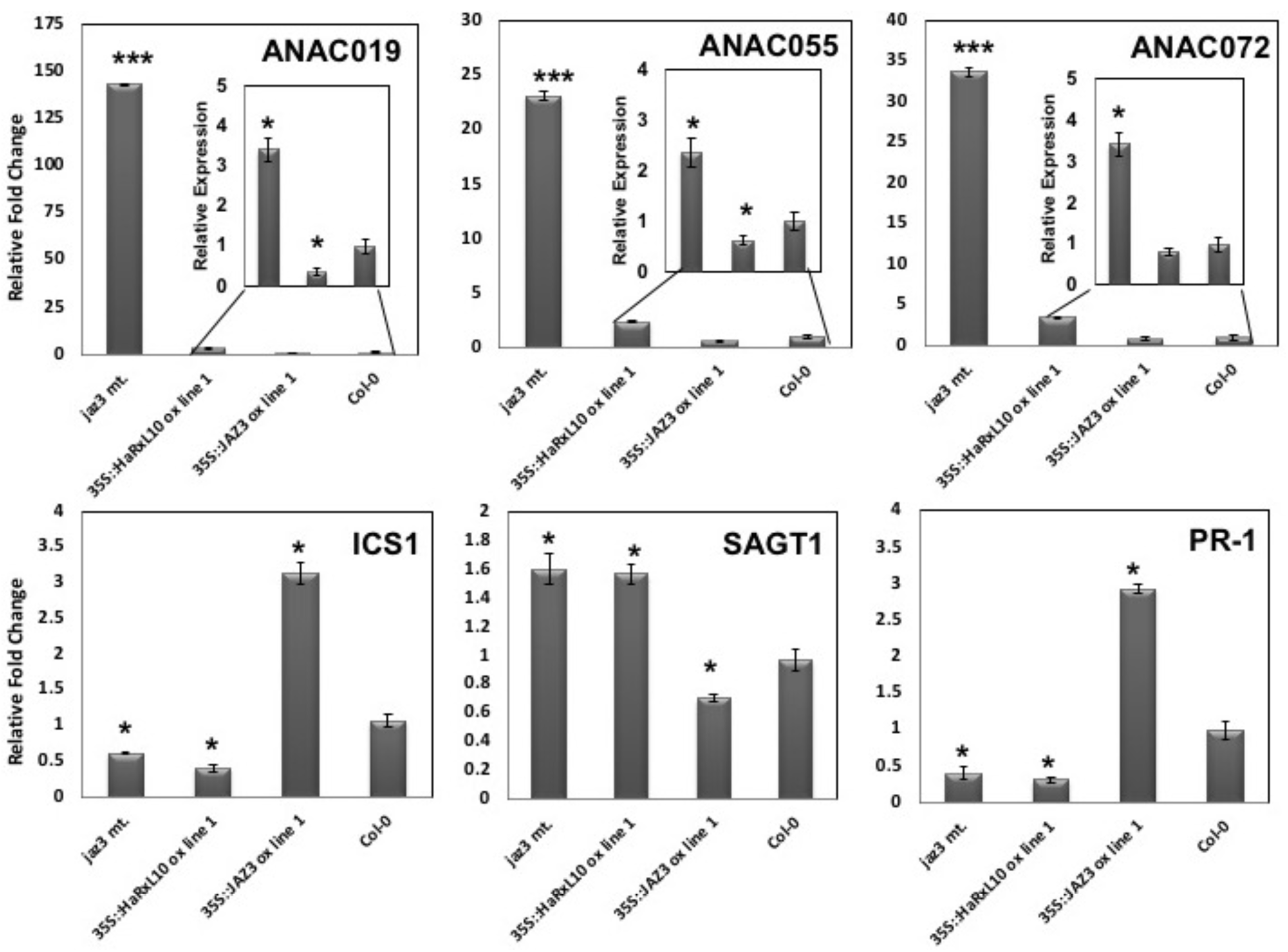
JAZ3 and HaRxL10 regulate expression of genes associated with SA biosynthesis and metabolism. Transcript abundance was measured using quantitative, real-time PCR using cDNA from *Hpa*-infected *jaz3* knockout mutant, Col:35S-RxL10 OX line 1 and Col:35S-JAZ3 OX line 1 with primers specific for the indicated genes. Samples were collected 24 hours post infection. Transcript abundance was normalized to *AtACTIN2*. Error bars depict variance among technical replicates. * ddCt values representing statistically significant (\**P* < 0.05) differences between transgenic or mutant lines and Col-0. This experiment was repeated three times with similar results.

Importantly, the *jaz3* knockout mutant was similar to Col:*35S-HaRxL10* in its effects on all six SA-associated genes (Fig. 8). Moreover, overexpression of *JAZ3* had the opposite effect: transcript abundance of *ANAC019*, *ANAC055*, *ANAC072*, and *SAGT1* is reduced in Col:35S-JAZ3, while *PR-1* and *ICS1* transcripts are elevated (Fig. 8). Together, these results establish an important role for JAZ3 in regulating the JA-SA crosstalk, and demonstrate that, remarkably, *HaRxL10* phenocopies the downstream effects of COR, and of a *jaz3* knockout, on the SA suppression cascade.

## DISCUSSION

Pathogens face a considerable challenge in avoiding or supressing the host plant’s immune responses. The plant immune signalling network is underpinned by substantial “cognitive diversity” (Dangl, 1993) provided by hundreds of immune surveillance proteins, and by a regulatory network comprised of signalling sectors that are genetically separable but interact in redundant, compensatory, or antagonistic configurations contingent on the nature of the attack with which the plant is faced (Katagiri & Tsuda, 2010; Nobori et al., 2018). This robust, adaptable immune system imposes intense selective pressure on pathogens to exploit points of vulnerability in the network. Accordingly, diverse pathogens and pests have independently evolved mechanisms through which phytohormone signalling within the immune network can be manipulated (Kemal Kazan & Lyons, 2014). This is best exemplified by co-option of JA-SA antagonistic cross talk: while this antagonism provides the plant with capacity for optimal defence, this configuration also carries a vulnerability, because pathogens/pests can activate one sector to suppress the other (Gutjahr & Paszkowski, 2009; Zhang et al., 2017). Understanding the mechanisms underpinning this co-option, across pathogen taxa, comprises an important piece in the puzzle of how microbes and pests successfully exploit host plants (Michelmore et al., 2017). In this report, we provide evidence for a novel mechanism through which an oomycete can manipulate JA-SA antagonism: HaRxL10 is secreted by *Hpa* into Arabidopsis cells, wherein it traffics to the nucleus and directly engages the JAZ3 transcriptional repressor protein to directly or indirectly trigger its degradation by the 26S proteasome. This activates a signalling cascade, involving transcriptional regulation of and by three NAC genes, that attenuates signalling through the SA sector (Figure 3-figure supplement 1). Attenuation of SA signalling is advantageous for *Hpa*, because SA-dependent signalling is a key aspect of immunity against biotrophic oomycetes. Treatment with exogenous SA is sufficient to induce resistance against oomycetes, and experiments with SA biosynthesis and signalling mutants have demonstrated that SA is often genetically necessary for NTI against avirulent oomycete strains, for basal resistance that limits the growth of virulent strains, and for systemic acquired resistance to limit the spread of the pathogen (Klessig et al., 2018; McDowell & Dangl, 2000). Mechanistically, SA signalling has been linked to programmed cell death which is a decisive defence against biotrophic pathogens, and with activation of *PR-1* for which an oomycete-specific resistance mechanism in sterol sequestration was recently described (Gamir et al., 2017).

For these reasons, it is likely that biotrophic and hemi-biotrophic oomycetes have evolved multiple mechanisms to interfere with SA-mediated signalling. This is supported by demonstrations that *Hpa* can suppress activation of *PR-1* specifically in plant cells containing haustoria, and by analysis of RXLR effectors that suppress SA-mediated responses when overexpressed (Asai et al., 2014; Caillaud et al., 2013). Now, mechanistic connections are beginning to emerge: One provided here by HaRxL10, and another from previous work on the *Hpa* RxLR effector, HaRxL44. Like HaRxL10, HaRxL44 targets regulatory cascades that act downstream of JA biosynthesis. However, HaRxL44 disrupts JA regulation at a different stage in the signalling hierarchy: HaRxL44 targets the MED19a subunit of the mediator transcriptional complex that recruits general transcription factors to plant gene promoters (Caillaud et al., 2013). This occurs downstream of SCF^COI1^-JAZ to activate MYC2 and other JA-responsive transcriptional complexes. As with HaRxL10 and JAZ3, HaRxL44 destabilizes MED19a by a proteasome-dependent mechanism. This manipulation of MED19a perturbs the Mediator complex to “shift the balance of defence transcription” (Caillaud et al., 2013) from SA-dependent immunity towards gene expression changes that typify responses to JA and ethylene. It is not currently clear whether the downstream responses include activation of the same MYC-*At*NAC cascade triggered by COR and HaRxL10, but the net effects are similar: SA signaling is compromised and plants are rendered more susceptible to *Hpa*. Thus, *Hpa* manipulates the JA pathway through at least two different mechanisms, one targeting the Mediator complex, and the other targeting a specific JAZ protein.

Strikingly, JAZ proteins are also targeted by effectors from a pathogenic bacterium and a mutualistic fungus (Gimenez-Ibanez, Chini, & Solano, 2016). Each of these effectors impinges on different sets of JAZ proteins through distinct mechanisms. The best-understood mechanism involves coronatine from *P. syringae*, which is a structural and functional mimic of jasmonic acid. In fact, COR is actually a more potent activator of the SCF^COI1^-JAZ receptor complex than is JA-Ile itself (Zhang et al., 2017). Thus, COR functions as a potent agonist of the SCF^COI1^-JAZ ubiquitin ligase complex, resulting in the degradation of most or all JAZ proteins. *P. syringae* can also deploy Type III effectors HopZ1a and HopX1 that target JAZ proteins for degradation (Gimenez-Ibanez et al., 2014; S. Jiang et al., 2013): HOPZ1a interacts with *Gm*JAZ1 from soybean and *At*JAZ, 2, 5, 6, 8, & 12 from Arabidopsis. *At*JAZ6 is acetylated by HOPZ1a and its degradation is dependent on COI1. *P. syringae* HOPX1 has a somewhat different mode of action: This effector is a cysteine protease that directly degrades the JAZ family as a whole, independently of COI1. Another *P. syringae* effector, HopBB1, exploits the SCF^COI1^-JAZ receptor complex in a different way: HopBB1 functions as an adaptor to link the Arabidopsis TCP14 transcription factor to JAZ3, thereby connecting TCP14 to SCF^COI1^ and triggering its degradation by the proteasome (Yang et al., 2017).

Additionally, HopBB1 disrupts the inhibitory interaction of JAZ3 with MYC2, leading to the activation of MYC2-regulated genes. Finally, the effector MiSSP7, from the mutualistic fungus *Laccaria bicolor*, has a completely different mode of action compared to bacterial effectors: In the interaction between *L. bicolor* and poplar roots, this protein is secreted in arbuscules to stabilize the poplar JAZ6 protein (Plett et al., 2014). This represses JA-responsive genes, thereby inhibiting JA-induced defence responses that would otherwise restrict fungal colonization (Plett et al., 2014). These studies, along with our data on HaRxL10, illustrate that the SCF^COI1^-JAZ regulatory hub is an Achilles Heel for diverse plants, subject to co-option by evolutionarily diverse pathogens that have independently evolved different mechanisms to manipulate JAZ proteins to their advantage.

In this context, it is interesting to compare HaRxL10-dependent effects to those from *P. syringae* effectors. First, HaRxL10 appears to be relatively specific: while COR, HopZ1a, or HopX1 degrade most or all of the JAZ proteins, HaRxL10 binds only to JAZ3, JAZ4, and JAZ9 in yeast two-hybrid assays, and our genetic experiments indicate that the interaction with *jaz3* holds the highest relevance for suppression of basal resistance to *Hpa*. This apparent specificity is similar to that of *P. syringae* HopBB1. As with HaRxL10, HopBB1 predominantly targets JAZ3. Intriguingly, HopBB1’s selective activation of MYC2-regulated genes and derepression of TCP14-regulated genes collectively activate only 20% of JA-responsive genes (Yang et al., 2017). Thus, HopBB1 is hypothesized to provide a strategy through which bacteria can exploit JA-dependent repression of immunity while avoiding negative effects associated with complete activation of the JA transcriptome (Yang et al., 2017). Perhaps the selective targeting of a small set of JAZ proteins by HaR×10 achieves the same specificity for *Hpa*, especially considering that HaRxL10 and HopBB1 both interact with JAZ3. However, our current data supporting the selectivity of HaRxL10 are based mostly on yeast two-hybrid assays and are therefore insufficient to definitively rule out biologically relevant interactions between HaRxL10 and other members of the JAZ family. We intend to pursue this question with comprehensive tests of all *jaz* mutants and double/triple combinations, along with interaction tests in systems other than yeast. This will provide a clearer picture of the extent to which other JAZ proteins are targeted by HaRxL10 and will also reveal whether other *AtJAZ* genes are relevant to the interaction with *Hpa*.

Another important question for the future is the exact mechanism through which HaRxL10 destabilizes JAZ3. Our protein degradation experiments in *N. benthamiana* with the proteasome inhibitor MG132 suggest involvement of the 26S proteasome, but the HaRxL10 protein does not contain any sequence motifs characteristic of E3 ligases. Thus, we hypothesize that HaRxL10 is an adapter protein that recruits JAZ3 to an E3 ubiquitin ligase complex. The simplest model is that RxL10 functionally mimics JA and COR by recruiting JAZ proteins to SCF^COI1^. However, RxL10 and COI1 do not interact in a yeast two-hybrid assay (Fig. 3). Thus, the involvement of COI1 requires further investigation. Another potential mechanism for *Ha*RxL10-mediated degradation of JAZ3 involves *At*BOI1: This protein interacts with *Ha*RxL10 in yeast (PPIN1, Figure 2-figure supplement 1A) and is a RING E3 ligase that positively regulates resistance to the necrotrophic fungus *Botrytis cinerea* (T. Mengiste et al., 2010). Future experiments will test whether COI1 and/or BOI1 contribute to *Hpa*-induced turnover of JAZ3.

The Arabidopsis *JAZ* gene family is comprised of 12 members, suggestive of substantial potential for functional redundancy (Chini et al., 2016). The mechanisms that underpin general and specific roles for *JAZ* genes are only beginning to emerge, and a specific role in oomycete immunity has not been identified for any individual *JAZ* gene to date. Thus, one of the most striking findings of our work is *JAZ3’*s genetically unique role in resistance to *Hpa*. With respect to bacteria, *JAZ2* has been identified as a major contributor to guard cell closure in stomatal defence and appears to be the primary target of COR in guard cells (Gimenez-Ibanez et al., 2017). *JAZ5* and *JAZ10* cooperatively restrict COR-induced phyto-toxicity, and a *jaz5/jaz10* double mutant displays moderately enhanced susceptibility to *P. syringae* (de Torres Zabala et al., 2016). *jaz3* mutants do not compromise stomatal defence against *P. syringae* (Gimenez-Ibanez et al., 2017) or display enhanced susceptibility to virulent *P. syringae* (de Torres Zabala et al., 2016), but they do partially suppress NTI against avirulent *P. syringae* (Mukhtar et al., 2011). The attributes of *JAZ3* that render it functionally unique in the interaction with *Hpa* remain to be determined and will be a major point of emphasis for future studies. Interestingly, two members of the YABBY transcription factor family appear to be targeted solely by JAZ3 and contribute to defence against *P. syringe* (Boter et al., 2015). These genes, along with *MYC2/3/4* and *ANAC19/55/072*, provide attractive leads for a more detailed understanding of how *JAZ3* mediates oomycete resistance.

In conclusion, our study of the *Hpa* effector HaRxL10 has enabled identification of a new mechanism through which an oomycete pathogen manipulates antagonistic JA-SA crosstalk to suppress a key immune signaling sector. Another notable aspect of this work is that it illustrates how studies of pathogen effector targets can provide new insights into the function of plant regulatory networks (Win et al., 2012): Just as studies of coronatine led to key insights into the JA receptor complex (Xie, Feys, James, Nieto-Rostro, & Turner, 1998), HaRxL10 has led us to the identification of *JAZ3* as an important gene for oomycete susceptibility and cross-talk between the JA and SA sectors. Finally, this work generalizes further that the JAZ signalling hub is a central target for virulence strategies of diverse prokaryotic and eukaryotic microbes that are beneficial or detrimental to their hosts. Collectively, our work and others’ lay out a clear road map of additional experimentation to better define important aspects of the JAZ hub’s functionality and its molecular characteristics that render it vulnerable to effector-mediated manipulation.

One potential payoff from such research lies in developing alleles of key JAZ proteins that are resistant to pathogen manipulation, as discussed in (Zhang et al., 2017). The HaRXL10-JAZ3 interaction is a promising test case for this approach, because HaRxL10 interacts only with a small number of JAZ proteins, one of which (JAZ3) has a clear pathogen phenotype with no redundant effects from other JAZ proteins. Importantly, to our knowledge no other unique functions have been ascribed to JAZ3 other than possible roles in salt stress (Valenzuela et al., 2016) along with JA-induced anthocyanin and chlorophyll loss (Boter et al., 2015). Thus, pleiotropic tradeoffs of JAZ3 mutation could be minimal. We intend to pursue these proof-of-concept experiments in Arabidopsis and to generalize the importance of the JAZ hub to other oomycete pathosystems.

## MATERIALS AND METHODS

### Construction of expression plasmids

HaRxL10:pDONR207 was kindly provided by B. M. Tyler. This clone was generated without the stop codon using standard Gateway cloning procedures (Invitrogen), and shuttled into expression plasmids using LR recombinase. For *Agrobacterium*-mediated transient assays and subcellular localization studies, HaRxL10-pDONR207 was shuttled into pB2GW7 and pEarleyGate104 vectors, respectively. For experiments involving *Pseudomonas syringae*, the entry construct of HaRxL10 were recombined into pEDV6 (Sohn et al., 2007). The expression plasmids obtained were mobilized from *E. coli* DH5α to coronatine-deficient mutant *Pst* DC3118 and effector-deletion mutant *Pst* DC3000 ΔCEL by standard triparental mating using *E. coli* pRK600 as a helper strain. For BiFC experiments, the pDONR207 clone of HaRxL10 was recombined into the pE-SPYNE-GW binary vector, which fuses the N-terminal half of YFP (nYFP) to the N-terminus of HaRxL10. Similarly, the pENTR4 construct of JAZ3 was cloned into the pE-SPYCE-GW binary vector, which fuses the C-terminal half of YFP (cYFP) to the N-terminus of JAZ3. The resulting binary expression plasmids were transformed into *Agrobacterium* GV3101. For *in vitro* co-IP studies, the entry clones of HaRxL10 and JAZ3 were recombined into the pIX-HA and pIX-GST binary vectors. For yeast-two-hybrid experiments, the *JAZ* and *COI1* coding sequences (CDS) were amplified using RT-PCR from total RNA extracted from *Arabidopsis thaliana* Col-0 and cloned into pCR2.1 (Life Technologies, Grand Island, NY). The coding sequences of the *JAZ* and *COI1* genes were released from plasmid pCR2.1 by digestion with *BamHI* and *EcoRI* restriction enzymes and the resulting fragments separated by agarose gel electrophoresis. The DNA fragments were purified using a Qiagen gel extraction kit (Qiagen, Valencia, CA). The *JAZ* coding sequences were ligated into the multi-cloning site of the Y2H vector pB42AD (Clontech, Mountain View, CA) to generate N-terminal fusions to the B42 transcriptional activation domain. The *RxL10* CDS was recombined from pDONR207 into a Gateway compatible version of the Y2H vector pGilda (Clontech, Mountainview, CA) to generate an N-terminal fusion to the LexA DNA binding domain. The Y2H constructs were transformed into *E. coli* DH5 alpha chemically competent cells and selected on LB plates with ampicillin. All clones were verified by sequencing.

### Plant growth conditions and generation of transgenic *Arabidopsis* plants

*Arabidopsis thaliana* and *Nicotiana benthamiana* plants were grown in Sunshine Mix #1 for all experiments. For pathogen experiments, *Arabidopsis* was grown under short day conditions (8 hours (h) light at 22°C and 16 h dark at 20°C. For all other experiments, *Arabidopsis* and *N. benthamiana* were grown at 16 h light and 8 h dark at 22°C. *Arabidopsis* Col-0 were transformed following the floral dip method (Clough & Bent, 1998). Primary transformed plants were selected on the basis of BASTA-resistance. The presence of transgenes and transcript abundance were confirmed by PCR from genomic DNA and qRT-PCR, respectively. Segregation assays were performed in the T2 generation to identify lines with a single transgene locus. Homozygous T3 or T4 plants were used in all experiments.

### *Hyaloperonospora arabidopsidis* maintenance, infection, and growth assays

Weekly propagation and maintenance of *Hyaloperonospora arabidopsidis* Emco5 was performed on the susceptible *Arabidopsis* ecotype, Ws-0, as described (McDowell, Hoff, Anderson, & Deegan, 2011). For *Hpa* growth assays, 10-12 day old *Arabidopsis* seedlings were sprayed with suspensions of 5×10^4^ sporangia/ml. Sporangiophore counts and trypan blue staining to visualize areas of cell death was performed as described (McDowell et al., 2011).

### RNA isolation, RT-PCR and qRT-PCR

Total RNA was isolated from uninfected and *Hpa* infected *Arabidopsis* seedlings using an RNeasy mini kit (Qiagen). To obtain cDNA for reverse-transcription and qRT-PCR, total RNA was first treated with DNase I (Ambion) and the first strand cDNA synthesis was performed using the OmniScript cDNA synthesis kit (Qiagen). Two micrograms of RNA were used as starting template material for the cDNA synthesis. 1µl cDNA was used per well with SYBR Green PCR Master Mix (Applied Biosystems) in 25µl reactions. Each PCR was performed in triplicate on the ABI7500 Real-time PCR system and transcript abundance was normalized to *AtACTIN2*.The primers used to detect specific transcripts are listed in Table S1. Statistical analyses were performed using Student’s t-test (*p < 0.05, **p < 0.01, ***p <0.001).

### Real Time PCR assay for growth of *H. arabidopsidis*

This assay followed the procedure described in (Anderson & McDowell, 2015). Briefly, five individuals from each sample were pooled in extraction buffer (200mM Tris pH 7.5, 25mM EDTA, pH 7.5, 250mM NaCl, 0.5% SDS) and genomic DNA (gDNA) was extracted using a bead beater. gDNA samples were quantified and diluted to 10 ng/uL final concentration. 25 µL samples were prepared by mixing 5 µL of gDNA sample with 12.5 µL of Sybr Green Mastermix (ABI, Carlsbad, California) along with primers and water. The primer sets *At*Actin Fwd/*At*Actin Rev were used for *AtACTIN* and *Ha*Act Fwd/*Ha*Act Rev were used for *HpaACTIN* (Brouwer et al., 2003). PCR reactions were performed on an ABI 7500 device. Ct values were determined using ABI software. Relative abundance to *AtACTIN* was calculated as 2^-dCt.

### *Pseudomonas syringae* infection

For spray inoculation assays, 4 to 5-week-old *Arabidopsis* plants were sprayed with 1×10^8^ cfu/ml bacterial solution in 10mM MgSO_4_ with 0.02% Silwet L-77. For syringe infiltration assays, 4 to 5-week-old plants were infiltrated using a needleless syringe with 1×10^5^ cfu/ml bacterial solution in 10mM MgSO_4_. Six plants were assayed for each data point. Leaf discs were cored at 0 days post infection (dpi) and 3dpi, surface sterilized with 70% ethanol and homogenized using a mini-bead beater (Biospec products). Serial dilutions were performed to count colony forming units. For each sample, three leaf discs were pooled three times per data point. Bacterial growth was measured as described previously Statistical analyses were performed on means of log-transformed data using Student’s t-test (*p < 0.05, **p < 0.01, ***p <0.001).

### Transient assays using *Agrobacterium*-infiltration in *N. benthamiana*

Recombinant *Agrobacterium tumefaciens* GV3101 were grown as described previously (Van der Hoorn et al., 2000) with the appropriate antibiotics. *Agrobacterium* liquid cultures were grown overnight, centrifuged, and resuspended in MMA induction buffer (10mM MgCl_2_, 10mM MES, 200mM Acetosyringone). The bacterial suspensions were incubated at room temperature for 1-3 hours. Infiltration using needleless syringe was performed on the abaxial side of 3-5 weeks old, *N. benthamiana* leaves. *Agrobacterium* strain containing pJL3-p19, a binary vector that expresses the suppressor of post-transcriptional gene silencing p19 of tomato bushy stunt virus (TBSV) (Voinnet, Rivas, Mestre, & Baulcombe, 2003) was co-infiltrated with the transformed *Agrobacterium* strains for enhanced expression. *Agrobacterium* strains carrying the respective constructs were mixed in 1:1:1 ratio along with pJL3-p19 in MMA induction buffer to a final OD_600_ of 0.3 (for confocal microscopy) and 0.5 (for western blotting). For co-expression experiments, *Agrobacterium* carrying YFP tagged-JAZ3 and 35S-HaRxL10 or 35S-HaRxL23 were mixed in 1:1 ratio along with pJL3-p19 in MMA induction buffer and syringe-infiltrated into 4 week-old *N. benthamiana* leaves. At 12 hours post infiltration, leaves were either syringe-infiltrated with water or 10 µM MeJA solution. Imaging was performed at 15 minute intervals after water or MeJA treatment. Imaging for BiFC experiments were performed 4-5 days post infiltration. All other imaging was performed 1-2 days post infiltration. Images were taken using confocal microscopy using a Zeiss Z.1, 25x or 40x water immersion objective and 488 HeNe laser. Processing of fluorescent images was performed using the Zeiss Zen 2012 software.

### Protein isolation and immunoblots

For western blotting, total proteins were extracted by grinding 3 to 4 leaf discs, each of 0.6 cm in diameter, in liquid nitrogen followed by boiling in SDS-loading buffer (50mM Tris-HCl, pH 7.5, 150mM NaCl, 1% Triton X-100, 0.1% SDS, 1mM EDTA, and 1mM DTT) with 1% protease inhibitor mix. Equal amounts of protein were separated on an SDS-polyacrylamide gel followed by semi-dry transfers onto nitrocellulose membrane (Whatman) using Hoefer SemiPhor apparatus for 30 minutes at 35-40mA. Membranes were blocked for four hours in 4% non-fat dry milk in TBS-T (50 mM Tris-HCl, pH 7.5, 150 mM NaCl, and 0.05% Tween 20). Overnight incubation at 4°C was performed with monoclonal anti-GFP antibodies (Covance Research) diluted with TBS-T (1:5000). After several washes with TBS-T the next day, the membrane was incubated with secondary anti-mouse Ig antibody (GE Healthcare) diluted with TBS-T for 1 hour at room temperature. The antibody-antigen complex was detected using HRP conjugated Immobilin Western Chemiluminescent substrate (Millipore). For JAZ3 degradation experiments in *N. benthamiana*, 3 to 4 leaf discs, each of 0.6 cm in diameter, were collected within 16 hours of *Agrobacterium*-infiltration for protein isolation and consecutive western blotting. For YFP-JAZ3 assays in *Arabidopsis*, 10 to 11 day old seedlings overexpressing YFP-JAZ3 were challenged with 50,000 spores/ml of *Hpa* Emco5. Tissues from infected and uninfected seedlings were collected, flash frozen in liquid nitrogen at the indicated time points and harvested later for protein isolation and western blotting as described above.

### *In-vitro* co-immunoprecipitation

*In vitro* pull-down assays were performed according to the manufacture’s protocol using the Pierce^®^ HA Tag IP/Co-IP Kit (Thermo Scientific). Briefly, HA- and GST-tagged proteins were synthesized *in vitro* using the TNT^®^ Coupled Wheat Germ Extract Systems (Promega). For the pull-down assays, equal amounts of N-terminal GST-tagged JAZ3 or GST alone and N-terminal HA-tagged HaRxL10 were incubated with gentle end-over-end mixing at 4°C with 20 µL anti-HA agarose slurry (35 µg antibody) overnight. Next day, the samples were washed 2 to 3 times with TBS-T (50 mM Tris-HCl, pH 7.5, 150 mM NaCl, and 0.05% Tween 20). After the final wash, the samples were subjected to SDS-PAGE followed by immunoblot analysis as described above using anti-GST antibody (Invitrogen).

### Yeast-2-hybrid screens

pGilda:HaRxL10 was co-transformed along with pB42AD:JAZ, pB42AD:COI1 or empty pB42AD constructs into yeast strain EGY48 carrying the p8Op:LacZ reporter plasmid. Yeast transformation reactions were selected on plates containing SD minimal media (BD Biosciences, San Jose, CA) supplemented with-uracil (U)/-tryptophan (W)/-histidine (H) amino acid drop out solution. Following selection, colonies were cultured overnight in liquid SD-UWH drop out media. The overnight cultures were harvested, washed 2X in sterile water, adjusted to OD_600_ = 0.2 and 10 µl of each culture was spotted onto agar plates containing SD galactose/raffinose-UWH media supplemented with X-gal (80 µg/ml). Y2H plates were incubated at 30 °C for 5 to 7 days and positive interactions/colonies were identified by development of blue color.

## ACKNOWLEDGEMENTS

Support was provided by the National Science Foundation Grant nos. IOS-0744875 and IOS-1353366 and by Agriculture and Food Research Initiative Competitive Grant no. 224426 from the USDA National Institute of Food and Agriculture. We thank Dr. Li Yang (Univ. of Georgia, USA) for the HaRxL44 construct and input on the experiments shown in Figure 4.

---

**Table S1:**
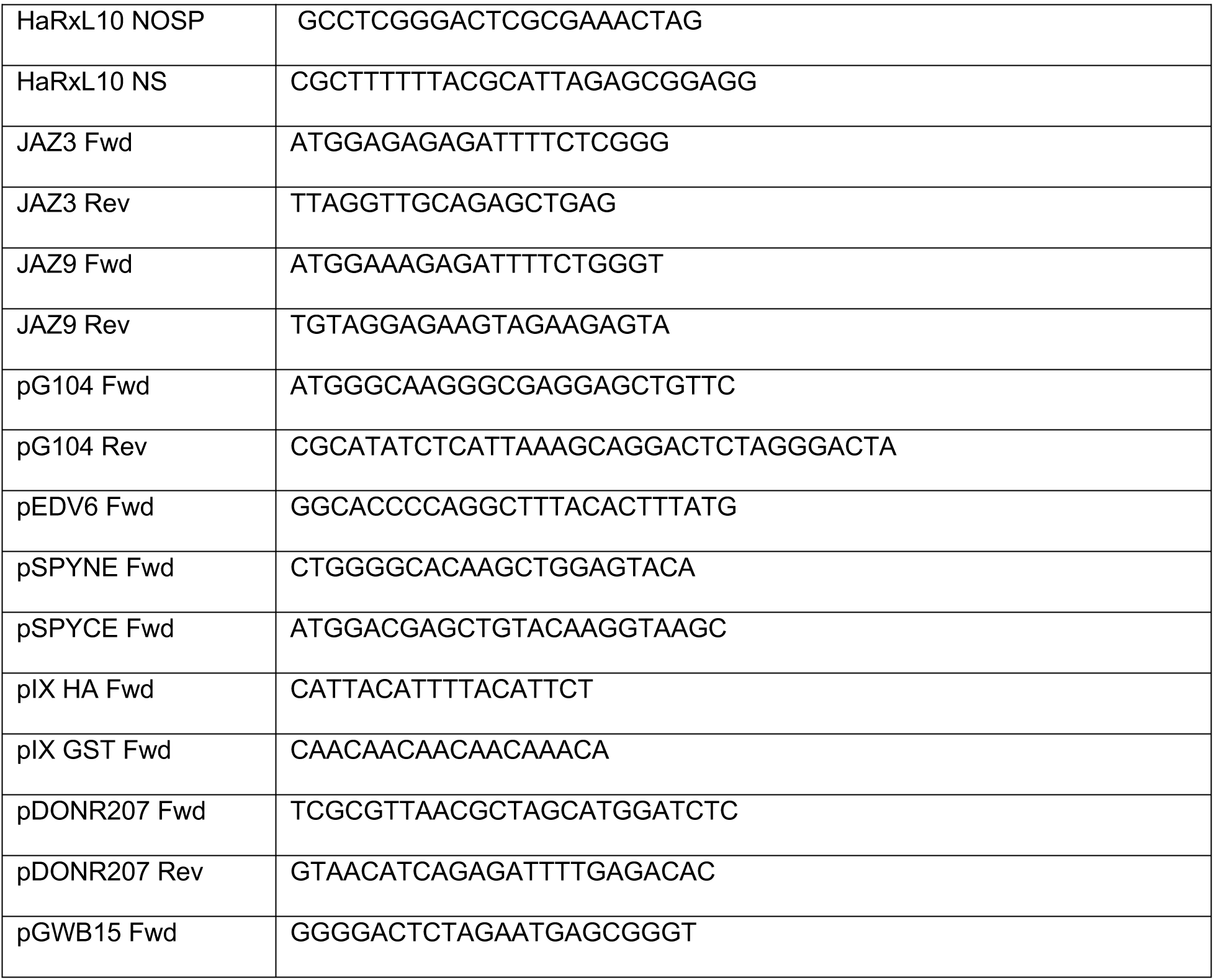
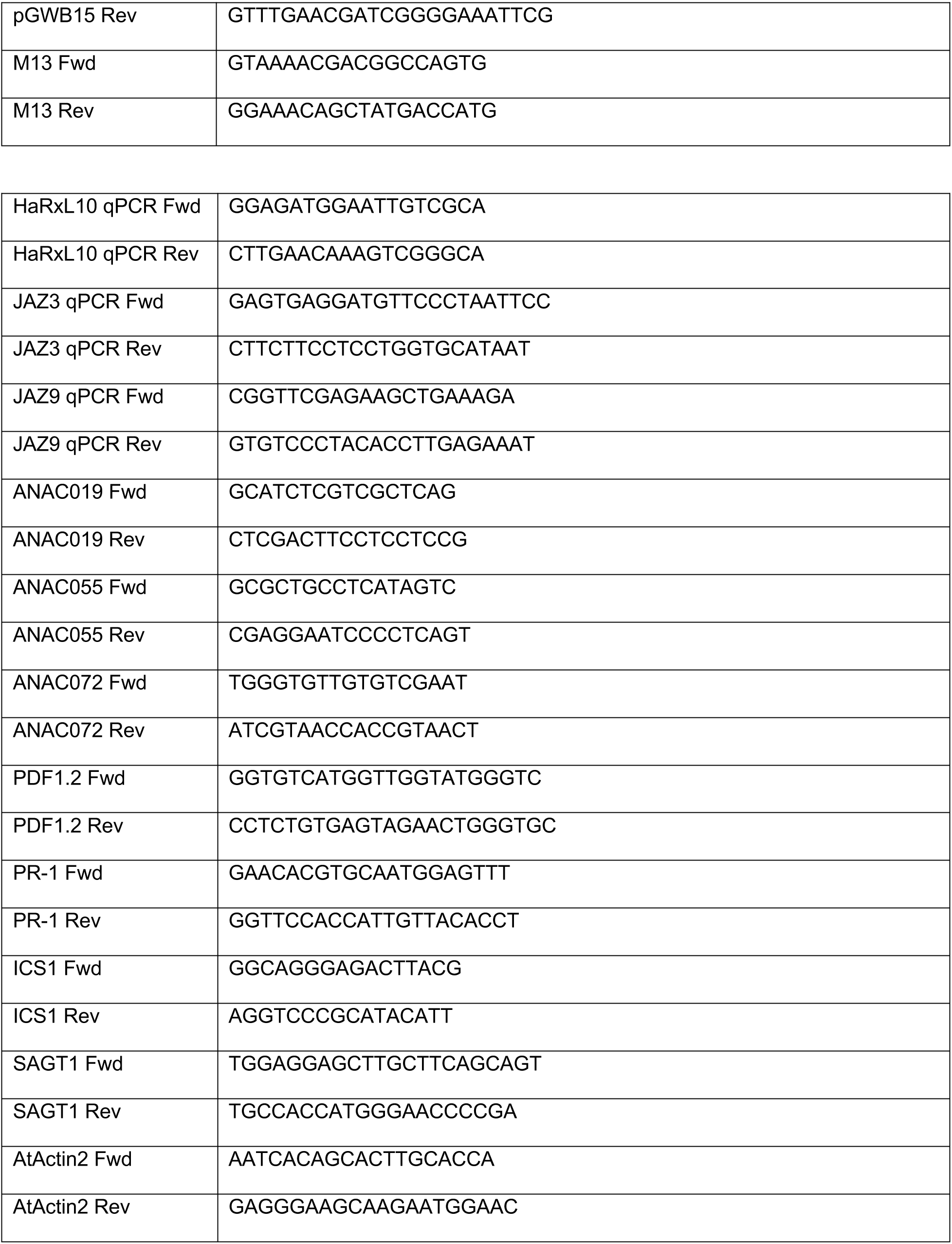
Oligonucleotide Primers

**Figure supplement 1.**
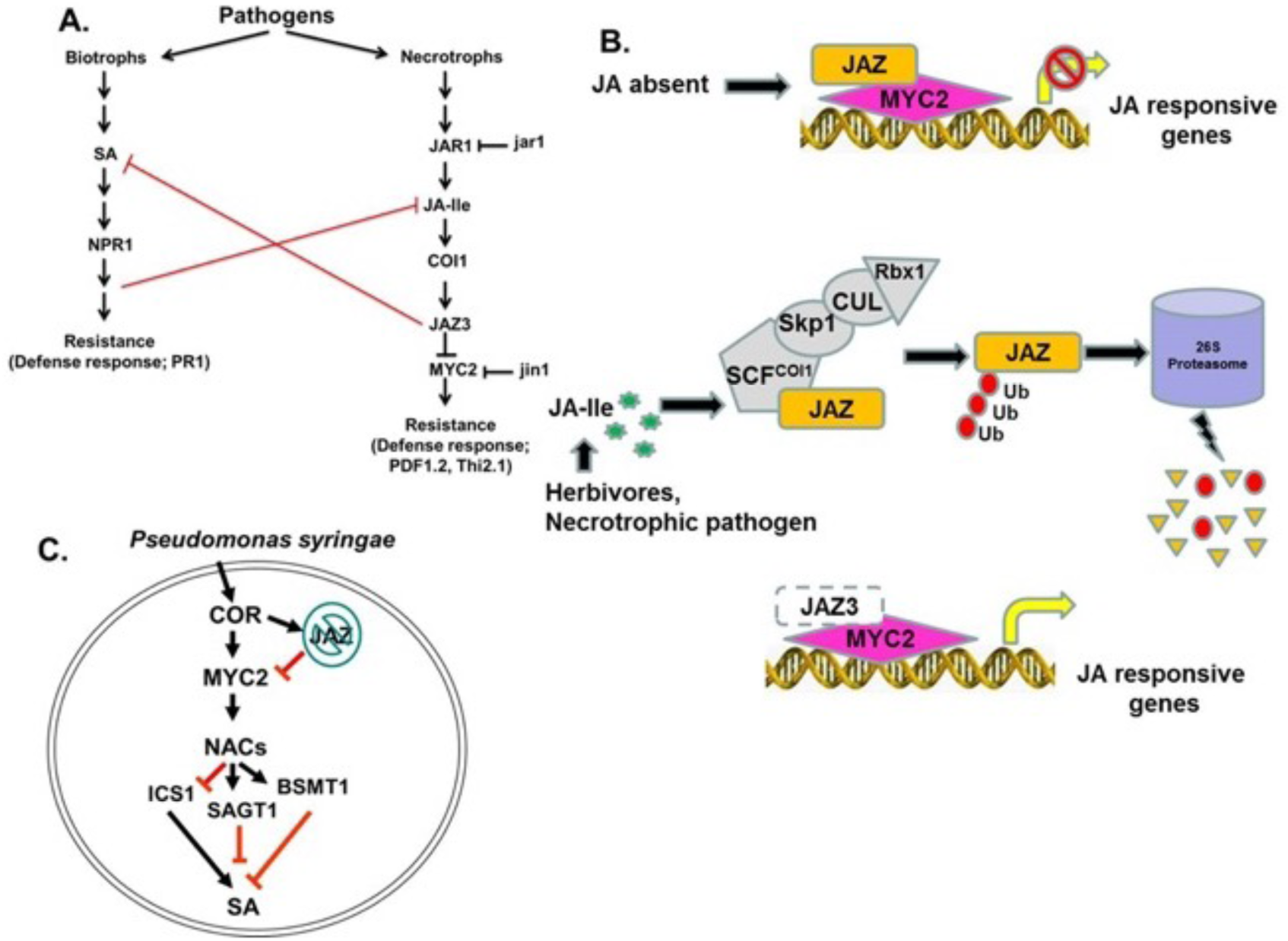
A schematic of JA biosynthesis, signalling, and manipulation by the bacterial toxin coronatine (COR). (**A**) Salicylic acid (SA) and jasmonic acid (JA) signaling sectors regulate resistance to biotrophic and necrotrophic pathogens, respectively, and are mutually antagonistic. Marker genes for each sector are indicated. (**B**) A model depicting major components of JA signalling. Absence of JA in plant cells enables repression of JA-responsive genes by JAZ proteins that repress the activity of transcription factors (e.g., MYC2). Repression is relieved by the initiation of developmental or environmental cues that increase the accumulation of bioactive JAs (JA-Ile). JA-Ile binds to and activates the SCF^COI1^ ubiquitin ligase complex. In turn, SCF^COI1^ targets JAZ proteins for ubiquitination and subsequent destruction in the 26S proteasome, thereby de-repressing downstream responses. (**C**) Mechanism for COR-induced suppression of SA accumulation in *Pseudomonas syringae*. COR acts as a molecular mimic of JA-Ile, thereby activating SCF^COI1^,which leads to degradation of JAZ proteins and activation of the NAC transcription factors through MYC2. NAC transcription factors then repress the SA biosynthesis gene *ICS1* and de-repress SA metabolism genes *BSMT1* and *SAGT1* to inhibit SA accumulation and promote bacterial virulence.

**Figure supplement 2.**
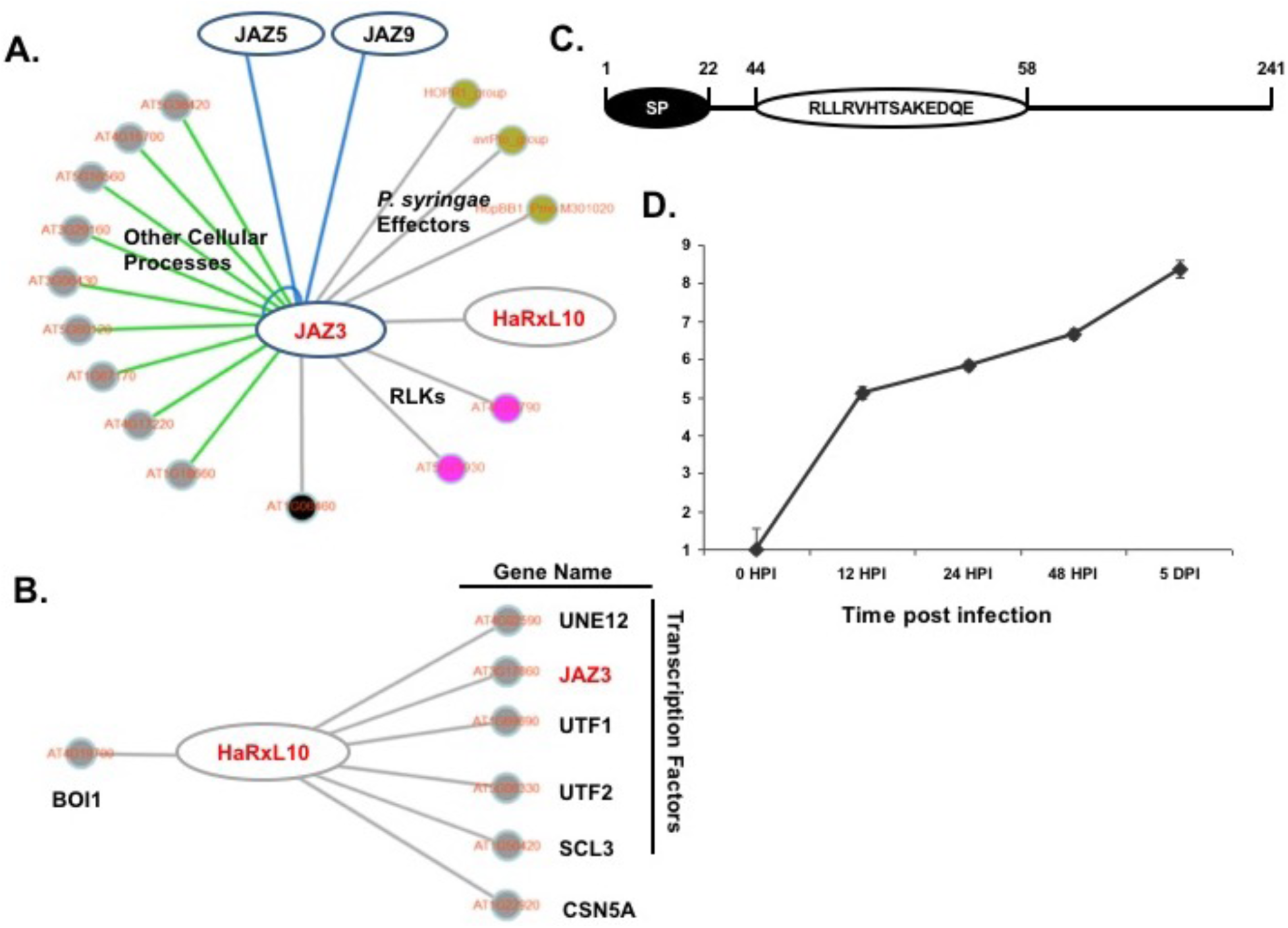
The *Hpa* effector HaRxL10 provides a potential connection to manipulation of JA responses. (**A**) Cytoscape schematic of proteins that interact with *Arabidopsis* JAZ3 in AtPPIN version 1. JAZ3 interacts with two other JAZ proteins, receptor like kinases (RLKs, pink), proteins involved in diverse cellular processes (grey), a defense related protein (black), three *P. syringae* effectors (gold), and *Hpa* effector HaRxL10. (**B**) Cytoscape schematic of proteins that interact with HaRxL10 in AtPPIN version 1. HaRxL10 interacts with four other transcriptional regulatory proteins, receptor like kinases (RLKs, pink), proteins involved in diverse cellular processes (grey), a defense related protein (black), three *P. syringae* effectors (gold), and *Hpa* effector HaRxL10. (**C**) Structural schematic of the HaRxL10 protein highlighting the signal peptide (SP) and the RXLR domain. (**D**) Transcript abundance of *HaRxL10* during infection by virulent *Hpa* Emco5. The abundance of *HaRxL10* was determined by qRT-PCR, using cDNA derived from Arabidopsis Col seedlings inoculated with virulent *Hpa* Emco5 and harvested at the indicated time points. The *x*-axis depicts transcript abundance of *HaRxL10* relative to *HpaActin*, normalized to expression at *t* = 0. Similar results were observed from three biological replicates.

**Figure supplement 3.**
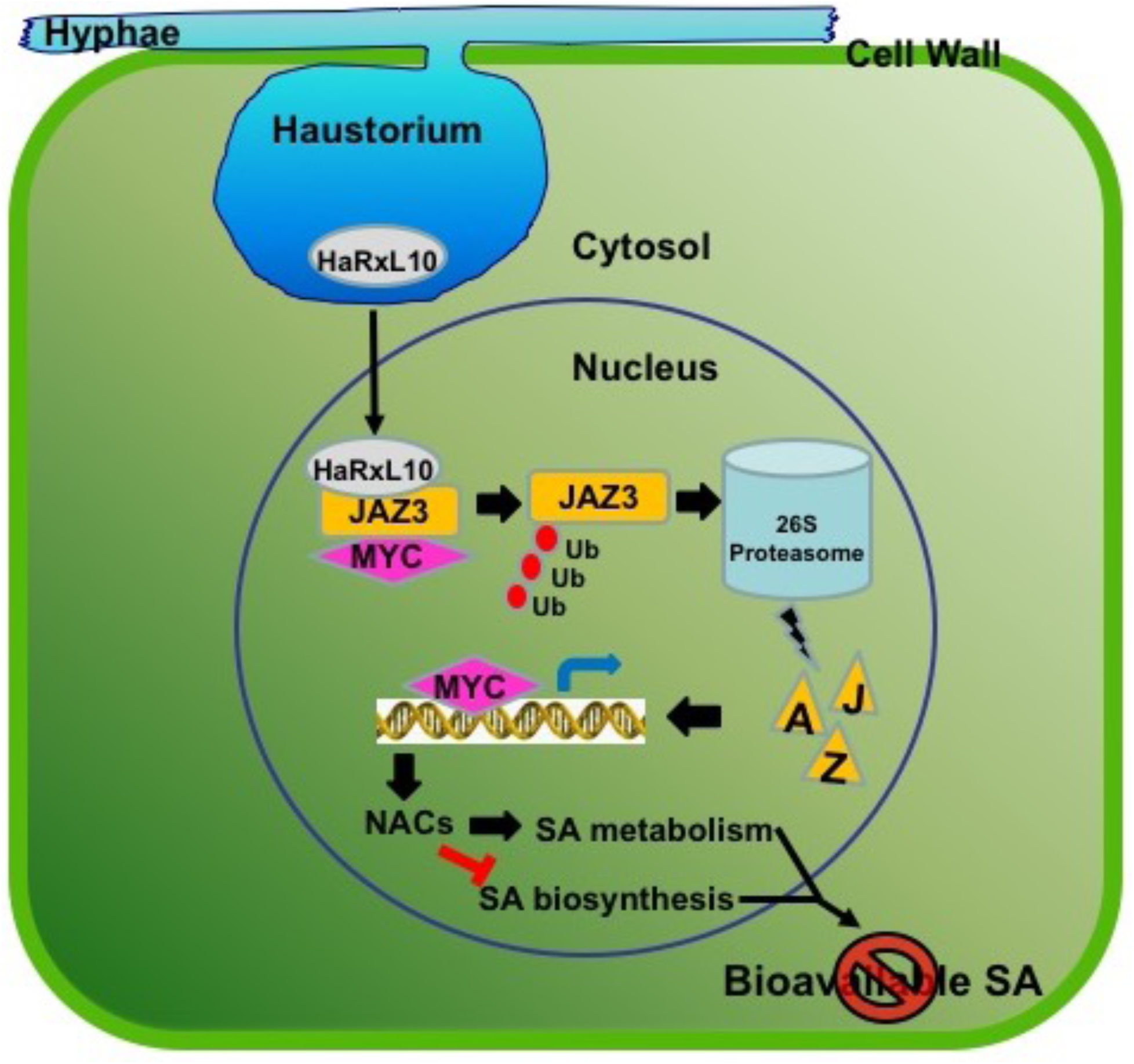
Hypothetical model showing the role of HaRxL10 in destabilizing JAZ3, thereby activating JA signalling and suppressing SA responses.

